# Spike-in probe-enhanced single-cell RNA-seq reveals post-infusion transcriptomic remodeling of “prime-and-kill” synNotch-CAR-T cells

**DOI:** 10.64898/2026.03.26.713760

**Authors:** Takahide Nejo, Payal B. Watchmaker, Milos S. Simic, Akane Yamamichi, Senthilnath Lakshmanachetty, Aishi Zhao, Jianwen Lu, Marco Gallus, Heather L. Benway, Robert Zhu, Ricardo Almeida, Wendell A. Lim, Hideho Okada

**Author notes:** **Corresponding author:** (H.O.). Pritzker School of Molecular Engineering, University of Chicago, Chicago, IL 60637, USA.

## Abstract

We previously developed synthetic Notch (synNotch)-chimeric antigen receptor (CAR)-T cells to improve the safety and efficacy of CAR-T therapy for glioblastoma. In this system, an anti-EphA2/IL13Rα2-dual-CAR is expressed only upon recognition of tumor- or brain-specific “priming” antigens, EGFRvIII (termed E-SYNC cells) or brevican (B-SYNC), respectively, with E-SYNC currently under phase I clinical evaluation (NCT06186401). However, tracking and profiling these engineered cells *in vivo* remain challenging, limiting our understanding of their activity and therapeutic potential. To address this gap, we developed a single-cell RNA-sequencing (scRNA-seq) workflow with custom spike-in probes for synNotch-CAR transcripts, enabling simultaneous detection of engineered cells and transcriptomic profiling. *In vitro*, integration of multiple probes using machine-learning-assisted classifiers detected 78.2% of E-SYNC cells and 60.0% of B-SYNC cells with 98.0% specificity. In a xenograft model, synNotch-positive cells were detected across the spleen, lung, and brain, with the highest frequency and most robust priming and activation observed in the brain. Single-cell transcriptomic analyses revealed tissue-specific differentiation programs, including cytotoxicity, proliferation, metabolic activity, and acquisition of tissue-resident memory phenotypes, shaped by both environmental cues and synNotch-mediated antigen recognition. In summary, this spike-in probe-enhanced scRNA-seq workflow enables robust detection and high-resolution characterization of synNotch-CAR-T cell dynamics and provides a broadly applicable platform for monitoring engineered immune cells in diverse clinical contexts.

**One Sentence Summary:** Our spike-in probe-enhanced single-cell RNA-sequencing method enables analysis of tissue-dependent activation and transcriptional states of synNotch-CAR-T cells, providing a robust and scalable platform for in vivo tracking and transcriptomic profiling of engineered cell therapies.

## Introduction

A major challenge for the successful development of chimeric antigen receptor (CAR)-T cell therapies for solid tumors is achieving reliable tumor recognition while avoiding off-target toxicity (*1*). In glioblastoma, the most common primary brain malignancy, there is a limited availability of antigens that are both tumor-specific and uniformly expressed across all cancer cells (*2*).

To address this challenge, we developed a synthetic Notch (synNotch)-CAR-T cell system, in which engineered T cells express a synNotch “priming” receptor that conditionally induces CAR expression upon recognition of a defined priming antigen. By selecting an appropriate priming antigen, this strategy enables the use of “killing” antigens that are more homogeneously expressed on tumor cells while minimizing the risk of off-target toxicity.

Specifically, we engineered synNotch receptors to recognize either the neoantigen epidermal growth factor receptor variant III (EGFRvIII) or brevican (BCAN), as glioblastoma-specific and central nervous system (CNS)-restricted priming antigens, respectively. Engagement of these priming antigens triggers expression of dual CAR reactive to ephrin type A receptor 2 (EphA2) and IL-13 receptor α2 (IL13Rα2) only within the brain microenvironment (*3–5*). In this system, CAR expression is spatially restricted, preventing on-target, off-tumor toxicity in non-brain tissues that may express EphA2 or IL13Rα2, as these killing antigens are normally absent in the brain. These anti-<u>E</u>GFRvIII or anti-<u>B</u>CAN <u>syn</u>Notch receptor-induced anti-EphA2/IL13Rα2 <u>C</u>AR-T cells are termed E-SYNC and B-SYNC, respectively. In both cases, synNotch-CAR-T cells achieved complete and durable clearance of patient-derived xenografts in murine glioblastoma models. Intravenously administered synNotch-CAR-T cells distribute systemically, but their activation is confined to tissues expressing the appropriate priming antigens (*3, 4*). Building on these promising preclinical results, we have initiated a phase 1 clinical trial evaluating E-SYNC cells (NCT06186401) (*6*).

Precise profiling of the biological kinetics and functional states of infused engineered immune cells is critical for understanding clinical responses, uncovering resistance mechanisms, and optimizing therapeutic strategies. Post-treatment patient samples from the clinical trial, such as blood and tumor tissues, provide a unique opportunity to validate this concept and uncover key therapeutic hallmarks of synNotch-CAR-T cells in humans.

However, tracking and profiling infused engineered cells *in vivo* remain challenging (*7*), particularly for complex cell therapies, such as synNotch-CAR systems. Clinical-grade E-SYNC and B-SYNC cells lack the fluorescent reporters present in their preclinical versions, and no reliable antibody-based assays currently exist for their detection. For example, the anti-G4S antibody used to detect specific CAR-T constructs (*8*) does not reliably detect E-SYNC or B-SYNC cells *in vivo*. Existing molecular approaches, including digital PCR (dPCR), RNAscope, and flow cytometry using fluorophore-conjugated proteins, provide useful information but have important limitations: dPCR lacks single-cell resolution, whereas RNAscope and flow cytometry do not enable comprehensive transcriptomic profiling.

Consequently, there is no robust method for sensitively detecting and functionally characterizing synNotch-CAR-T cells at single-cell resolution in clinical specimens.

Single-cell RNA sequencing (scRNA-seq) offers a potential solution (*9*). In published clinical trials, CAR-T cells have been profiled either after fluorescence-activated cell sorting (FACS)-based enrichment of CAR-expressing cells (*10–16*), by customizing the reference genome to computationally detect CAR transcripts (*17–24*), or through combinations of these approaches (*25, 26*). However, FACS-based strategies are largely restricted to conventional CAR constructs, such as anti-CD19 or anti-BCMA. Alignment-based detection often suffers from low transcript capture efficiency *in vivo* (*21*), sometimes requiring computational inference (*27, 28*). These sensitivity limitations are particularly problematic for synNotch-CAR-T cells, which may be present at low frequencies and exhibit variable transcript expression.

Here, we present a probe-based scRNA-seq strategy for robust detection and functional profiling of synNotch-CAR-T cells. Using the 10X Genomics Flex Gene Expression platform, we incorporated custom spike-in probes targeting synNotch-CAR transcripts into a premanufactured whole-transcriptome probe set. Applied to both *in vitro* conditions and *in vivo* xenograft samples, this approach enables robust detection of engineered cells while simultaneously capturing their transcriptomic states at single-cell resolution, thereby offering a scalable framework for clinical sample analysis.

## Results

### Designing custom spike-in probes for synNotch-CAR-T cell detection

To detect synNotch-CAR-T cells using probe-based scRNA-seq, we designed a panel of custom spike-in probes targeting specific sequences within three regions of the synNotch-CAR vectors: (1) the anti-EGFRvIII synNotch receptor (1,692 bp) in E-SYNC cells; (2) the anti-BCAN synNotch receptor (1,698 bp) in B-SYNC cells; and (3) the anti-EphA2/IL13Rα2-dual-CAR (1,173 bp), which is shared by both E-SYNC and B-SYNC cells (**Fig. 1**). The synNotch transcripts of both E-SYNC and B-SYNC contain a partial murine *Notch1* transgene and are constitutively expressed by design, whereas expression of the CAR transcript is inducible upon recognition of the appropriate priming antigen.

**Fig. 1.**
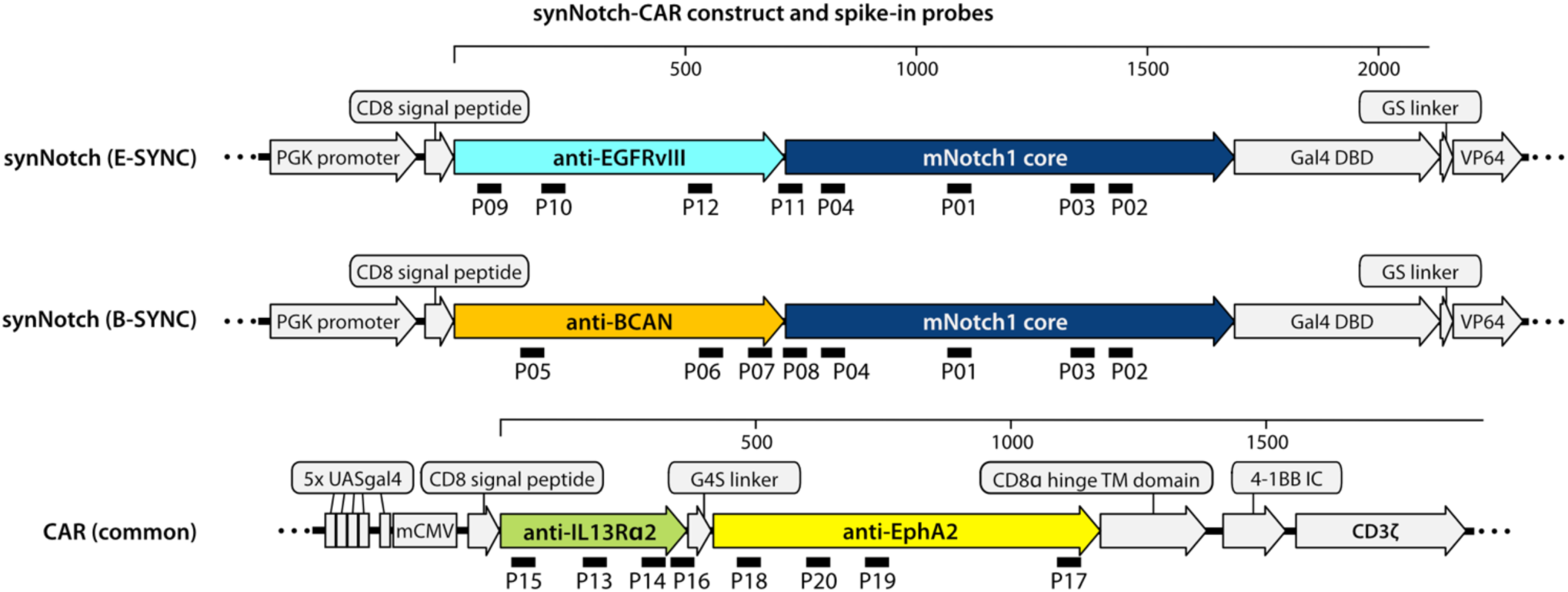
Designing custom spike-in probes for synNotch-CAR-T cell detection. Schematics of the constitutively expressed E-SYNC (**top**) and B-SYNC (**middle**) synNotch transcripts, and the common CAR transcript (**bottom**) expressed upon priming signals. Bars indicate regions targeted by custom spike-in probes. Details are provided in **Supplementary Table S1**. DBD, DNA-binding domain; UAS, upstream activation sequence. TM, transmembrane; IC, intracellular.

To select candidate probe sequences, we applied a systematic approach. First, we screened all possible 50-base sequences and retained those that met the recommended design criteria (e.g., a thymine at the 25^th^ position of each probe and GC content within a defined range) (*29*). Next, we performed a BLAST search against the human transcriptome to exclude candidates with low specificity. Of the 181 pass-filter sequences, we selected 45 candidates, evaluated their specificity using reverse transcription quantitative PCR (RT-qPCR), and established a final panel of 20 probes: four targeting the synNotch region shared by E-SYNC and B-SYNC (termed P01–P04), four specific to B-SYNC (P05–P08), four specific to E-SYNC (P09–P12), and eight targeting the CAR transcript (P13–P20) (**Fig. 1; Supplementary Table S1**). Probes P01–P12 target constitutively expressed synNotch transcripts for cell detection, while P13–P20 target the inducible CAR transcript to assess priming status.

### Samples and data processing for scRNA-seq

We performed a scRNA-seq experiment incorporating 20 custom spike-in probes across 16 samples (5 *in vitro* and 11 *in vivo*). The *in vitro* set comprised unprimed and primed E-SYNC cells, unprimed and primed B-SYNC cells, and untransduced (UT) human T cells (**Fig. 2A**; see **Methods** for details). E-SYNC and B-SYNC cells were generated from distinct healthy donors via lentiviral transduction using preclinical-grade dual lentiviral vectors carrying a Myc tag and fluorescent reporters (blue fluorescent protein [BFP] for synNotch and green fluorescent protein [GFP] for CAR). The UT sample was derived from the same donor as the B-SYNC cells. The use of preclinical-grade vectors enabled FACS sorting of BFP^+^/Myc^+^ double-positive cells to near-100% purity of synNotch-expressing cells, which is not achievable with clinical-grade vectors lacking fluorescent markers. Despite FACS purification to >99% purity, synNotch positivity consistently declined to 60–80% in both E-SYNC and B-SYNC cells by the time of *in vivo* administration or cell fixation, possibly due to transgene silencing (**Fig. S1A, B**). Primed E-SYNC and B-SYNC samples were prepared by 24-h stimulation with EGFRvIII beads or BCAN-coated culture plates, respectively (*3, 4*).

**Fig. 2.**
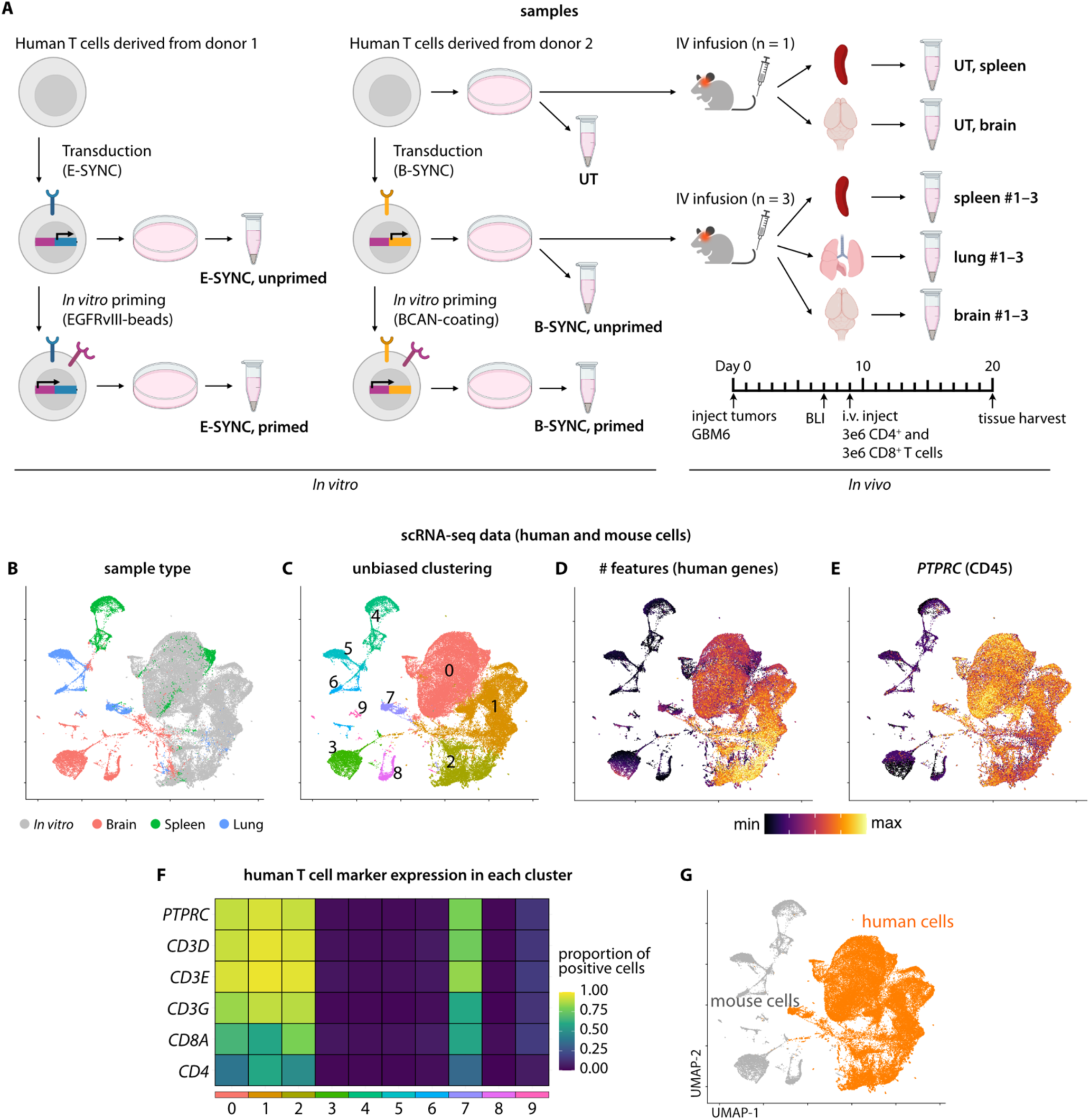
Overview of scRNA-seq experiment and detection of human cells. **(A)** Overview of the 16 samples analyzed. The same batches of unprimed B-SYNC and UT cells were intravenously administered to tumor-bearing mice. (**B–E**) Uniform Manifold Approximation and Projection (UMAP) plots showing sample types (**B**), 10 clusters identified by unbiased clustering (**C**), the number of features (human genes) per cell (**D**), and *PTPRC*/CD45 expression levels (**E**). (**F**) Heatmap showing the proportion of positive cells for human T cell marker genes in each cluster. (**G**) UMAP plot illustrating the distribution of human versus mouse cells based on expression profiles.

To investigate the *in vivo* dynamics of B-SYNC cells after infusion, we established an orthotopic xenograft model using the patient-derived glioblastoma cell line GBM6 (2 × 10^4^ cells), intracerebrally implanted into NOD CRISPR *Prkdc Il2rg* (NCG) triple-immunodeficient mice on day 0. Tumor engraftment was confirmed by bioluminescence imaging (BLI) on day 7. On day 9, mice received an intravenous infusion of unprimed B-SYNC or UT cells (3 × 10^6^ CD8^+^ and 3 × 10^6^ CD4^+^ cells per mouse). Mice were euthanized on day 20, and mononuclear cells were isolated from the spleen, lung, and brain for downstream analyses (**Fig. 2A**; see **Methods** for details).

In the resulting scRNA-seq dataset, we analyzed a total of 71,541 cells across the 16 samples (*in vitro*: mean 10,787; range 8,912–13,867; *in vivo*: mean 1,600; range 748–2,307). Additional quality control (QC) metrics are provided in **Supplementary Table S2**. All data were processed together using Seurat, including QC filtering, normalization, and scaling (**Fig. 2B**).

Unsupervised clustering identified 10 clusters (**Fig. 2C**), of which four (clusters 0, 1, 2, and 7) exhibited substantially higher human gene feature counts (nFeature_RNA values) and markedly increased expression of human T cell markers (*PTPRC*/CD45 and *CD3D*) (**Fig. 2D– F**). These four clusters comprised 58,455 cells and were therefore classified as human cells (*in vitro*: mean 10,787; range 8,912–13,867, all retained; *in vivo*: mean 411; range 2–818) (**Fig. 2G**).

Notably, in the brain-derived sample from the UT-treated mouse, only two human cells remained after filtering (**Supplementary Table S2**). This is consistent with our previous observation that, despite comparable *ex vivo* preparation methods, UT cells rarely migrate to or persist in the brain, whereas B-SYNC cells exhibit strong brain tropism, proliferation, and persistence (*4*). Accordingly, this sample was excluded from downstream analyses.

### Detection of synNotch-positive cells

Screening of the 20 spike-in probes revealed that 11 exhibited human-specific reactivity, whereas 3 E-SYNC-detecting probes (P09, P10, P12) produced negligible signals and were excluded from further analyses (**Fig. S2**). Six probes (P01–P04, P08, and P11) showed reactivity with both human and mouse cells, likely reflecting recognition of murine *Notch1* transcripts expressed endogenously in mouse cells or introduced as a transgene in synNotch-positive cells (**Fig. 1**). Importantly, while this cross-reactivity may affect mixed-species datasets, it does not impact analyses restricted to human cells (e.g., clinical samples); consequently, 17 spike-in probes were retained for downstream analyses.

We first evaluated nine individual spike-in synNotch–detecting probes (P01–P08 and P11) using five *in vitro* samples (n = 53,936 cells). E-SYNC and B-SYNC cells, both unprimed and primed, served as positive controls, while UT cells served as negative controls; detection was defined as a unique molecular identifier (UMI) count ≥ 1. For E-SYNC, the average detection rate across the five relevant probes (P01–P04 and P11) was 45.0% (range: 20.6–52.7%), whereas for B-SYNC, the average detection rate across the eight relevant probes (P01–P08) was 22.5% (range: 14.4–27.3%). Across all nine probes, the average specificity was 99.8% (range: 98.7–100%) (**Fig. S3**).

To enhance detection accuracy, we integrated signals across multiple probes, including eight CAR-targeting probes (yielding a total of 13 probes for E-SYNC and 16 for B-SYNC). For each cell, the maximum UMI count across probes (maxUMI_synNotch_) was calculated as a representative metric. An optimal threshold was determined using Youden’s J statistic to balance sensitivity and specificity (*30*), resulting in detection rates of 78.2% for E-SYNC and 60.0% for B-SYNC, with a specificity of 98.0% for both (**Fig. 3A; Fig. S4A–F**). We additionally trained machine learning (ML) classifiers using four algorithms: (1) generalized linear model (GLM; logistic regression), (2) elastic net-regularized logistic regression (GLMnet), (3) random forest (RF), and (4) support vector machine (SVM). Notably, all four ML classifiers showed complete concordance with the maxUMI-based classifier, supporting its robustness. Given the reported 20–40% frequency of synNotch gene silencing (**Fig. S1**), these detection rates correspond to capturing ≥95% of synNotch-positive E-SYNC cells and ∼72–96% of synNotch-positive B-SYNC cells.

**Fig. 3.**
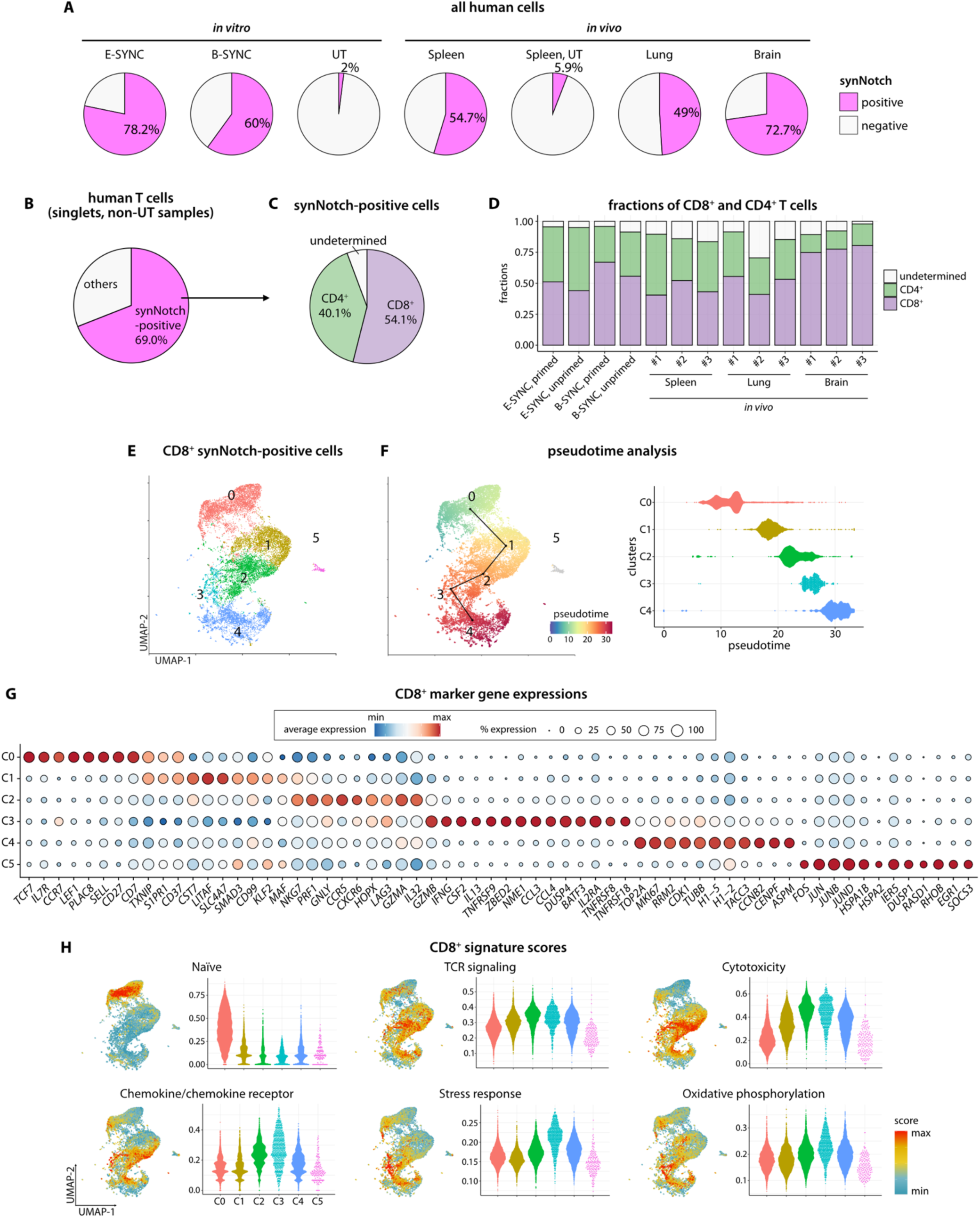
Detection and single-cell transcriptomic profiling of synNotch-positive cells. **(A)** Pie charts showing the proportions of human cells classified as synNotch-positive in each sample type, based on integrated signals from multiple probes. (**B–C**) Pie charts showing the proportion of synNotch-positive cells among singlet, non-UT human cells (**B**) and their distribution between CD8^+^ and CD4^+^ T cell subsets **(D)** Bar plot comparing the relative frequencies of CD8^+^ and CD4^+^ T cells across samples. **(E)** UMAP plot of synNotch-positive CD8^+^ cells colored by cluster identity. **(F)** UMAP feature plot and corresponding swarm plot showing pseudotime estimates (slingshot) for individual cells from clusters C0–C4. **(G)** Dot plot showing curated differentially expressed genes (DEGs) upregulated in each cluster. The complete DEG list is provided in **Supplementary Table S3**. **(H)** Swarm plots and UMAP feature plots illustrating representative CD8^+^ signature score distributions. The full set of 19 analyzed signatures is provided in **Fig. S6A**.

We next applied the *in vitro* B-SYNC-derived classification models to the *in vivo* dataset.

Using these models, 54.7%, 49.0%, and 72.7% of human T cells isolated from the spleen, lung, and brain of B-SYNC-treated mice, respectively, were classified as synNotch-positive (**Fig. 3A; Fig. S4G**).

A small degree of misclassification was observed in negative controls. In the spleen of the UT-treated mouse, seven cells (5.9%) were classified as synNotch-positive. Six of these were detected exclusively by probes P01–P04, which recognize the murine *Notch1* transgene (**Fig. S5A**). These cells were likely mouse cells erroneously classified as human, indicating that clustering-based discrimination between mouse and human cells is not entirely error-free. However, this issue is confined to mixed-species datasets. In contrast, among 163 cells (2.0%) from the *in vitro* UT sample classified as synNotch-positive, none were recurrently detected by multiple probes (**Fig. S5B**).

Collectively, these findings demonstrate that integrating signals across multiple probes enables robust detection of synNotch-positive cells in both *in vitro* and *in vivo* settings.

### Single-cell transcriptomic states of synNotch-positive cells

Having established robust detection of synNotch-positive cells, we characterized their transcriptomic states to define their functional heterogeneity and differentiation trajectories. Among 42,687 human cells from *in vitro* and *in vivo* non-UT samples, spike-in probe analysis identified 29,465 synNotch-positive cells (69.0%) (**Fig. 3B**). Of these, 54.1% were CD8^+^ and 40.1% were CD4^+^ T cells (**Fig. 3C**). Brain-derived samples contained a higher proportion of CD8^+^ T cells than other *in vivo* tissues (**Fig. 3D**), consistent with the reported proliferative advantage of CD8^+^ T cells driven by recognition of CAR antigens on tumor cells (*31*). Hereafter, CD8^+^ and CD4^+^ subsets were analyzed separately.

Within the CD8^+^ population, unsupervised clustering identified six clusters (C0–C5) (**Fig. 3E**). Pseudotime analysis placed C0 at the earliest state and C4 at the most differentiated state (**Fig. 3F**). Differential expression analysis characterized the clusters as follows: C0 comprised quiescent, naïve/central-memory-like cells enriched for *TCF7/*TCF-1, *LEF1*, *CCR7*,*SELL*/CD62L, and *IL7R*/CD127; C1 contained quiescent central-memory-like cells expressing *KLF2, TXNIP, and S1PR1*; C2 comprised cytotoxic effector cells with tissue-resident memory (TRM)-like features, characterized by strong cytotoxic programs (*GNLY, PRF1, GZMA,* and *NKG7*), trafficking molecules (*CCR5*, *CXCR6*, and *ADGRG1*/GPR56), tissue adaptation factors (*HOPX and IL32*), and inhibitory receptors (*LAG3* and *TIGIT*); C3 consisted of activated, polyfunctional effector cells expressing *CSF2*/GM-CSF, *IFNG*, *CCL3*, *CCL4*, and *BATF3*, and high levels of co-stimulatory molecules (*TNFRSF9*/4-1BB, *TNFRSF18*/GITR, and *TNFRSF8*/CD30); C4 represented proliferating cells enriched for *MKI67, TOP2A,* and *CDK1*; and C5 comprised rare, stress-responsive cells (1.1%, n = 183 cells) enriched for *FOS, JUN, JUNB, DUSP1, SOCS3, HSPA2,* and *HSPA1B* (**Fig. 3G; Supplementary Table S3**). Gene signature analysis confirmed enrichment of naïve-associated signatures in C0, whereas C2– C4 were enriched for T-cell receptor (TCR) signaling, cytotoxicity, and chemokine/chemokine receptor activity, with C3 showing particular enrichment for stress response, NFκB signaling, and oxidative phosphorylation (**Fig. 3H; Fig. S6A)**. Projection onto a reference human CD8^+^ tumor-infiltrating lymphocyte (TIL) dataset using ProjecTILs (*32, 33*) corroborated these assignments, with C0 aligning with naïve T cells, C1 with central- and effector-memory cells, and C2–C4 overlapping primarily with precursor- or terminally-exhausted populations (**Fig. S6B**).

In the CD4^+^ population, unsupervised clustering identified four clusters (C0–C3) (**Fig. S7A**), with pseudotime analysis placing C3 as the most differentiated state (**Fig. S7B**). Differential expression analysis characterized the clusters as follows: C0 comprised naïve/central-memory-like cells enriched for lymphoid-homing and stem-like signatures (*TCF7, LEF1, CCR7, SELL,* and *IL7R*); C1 contained central-memory-like cells with features suggestive of Th17 differentiation (*KLF2, S1PR1, S1PR4,* and *ITGB1*); C2 included activated effector cells with cytotoxic/Th1-like features (*GZMB*, *GNLY*, *CCL5*, and *IL2RB*/CD122), together with activation markers (*IL2RA*/CD25 and *TNFRSF4*/OX40) and inhibitory receptors associated with exhaustion or regulatory programs (*HAVCR2/*TIM-3, *TIGIT*, and *ENTPD1/*CD39); and C3 comprised proliferating cells enriched for cell-cycle genes (*MKI67, TOP2A,* and *CDK1*) and replication-linked histone genes (**Fig. S7C; Supplementary Table S4**). Gene signature analysis revealed enrichment of naïve-associated signatures in C0–C1; effector activation, TCR signaling, Treg-associated signatures, and costimulatory molecules in C2; and metabolic programs in C3 (**Fig. S7D**). Projection onto a reference CD4^+^ TIL dataset (*34*) using ProjecTILs corroborated these assignments: C0–C1 aligned with naïve T cells, C2 corresponded to cytotoxic T cells exhibiting features of exhaustion, and C3 represented a heterogeneous population that included regulatory T cells and Th17 helper T cells (**Fig. S7E**).

In summary, these analyses demonstrate that the probe-based scRNA-seq approach enables high-resolution profiling of transcriptomic states in synNotch-positive cells.

### Tracking the transcriptomic evolution of post-infusion synNotch-positive cells

To determine how tissue microenvironments shape synNotch-CAR-T cell differentiation after systemic infusion, we compared the pre-infusion sample (*in vitro* unprimed B-SYNC cells, representing the exact batch infused into the mice) with cells recovered from the spleen, lung, and brain 11 days post-administration.

Within the CD8^+^ population, clusters C0 and C1 predominated in pre-infusion (82.8% of cells), spleen (65.5%), and lung (60.5%) samples but were markedly reduced in the brain (31.9%), where clusters C2–C4 were the predominant populations (66.1%) (**Fig. 4A**). UMAP density plots confirmed this compositional shift: pre-infusion, spleen, and lung samples were enriched in naïve cells, whereas brain-derived cells were concentrated in activated and proliferating states (C2–C4) (**Fig. 4B**). Consistent with this shift, brain-derived cells exhibited elevated signatures of cytotoxicity, chemokine/chemokine receptor activity, exhaustion-associated programs, stress response, and multiple metabolic pathways, including oxidative phosphorylation, glycolysis, and fatty acid metabolism, compared with spleen and lung samples (**Fig. 4C; Fig. S8**).

**Fig. 4.**
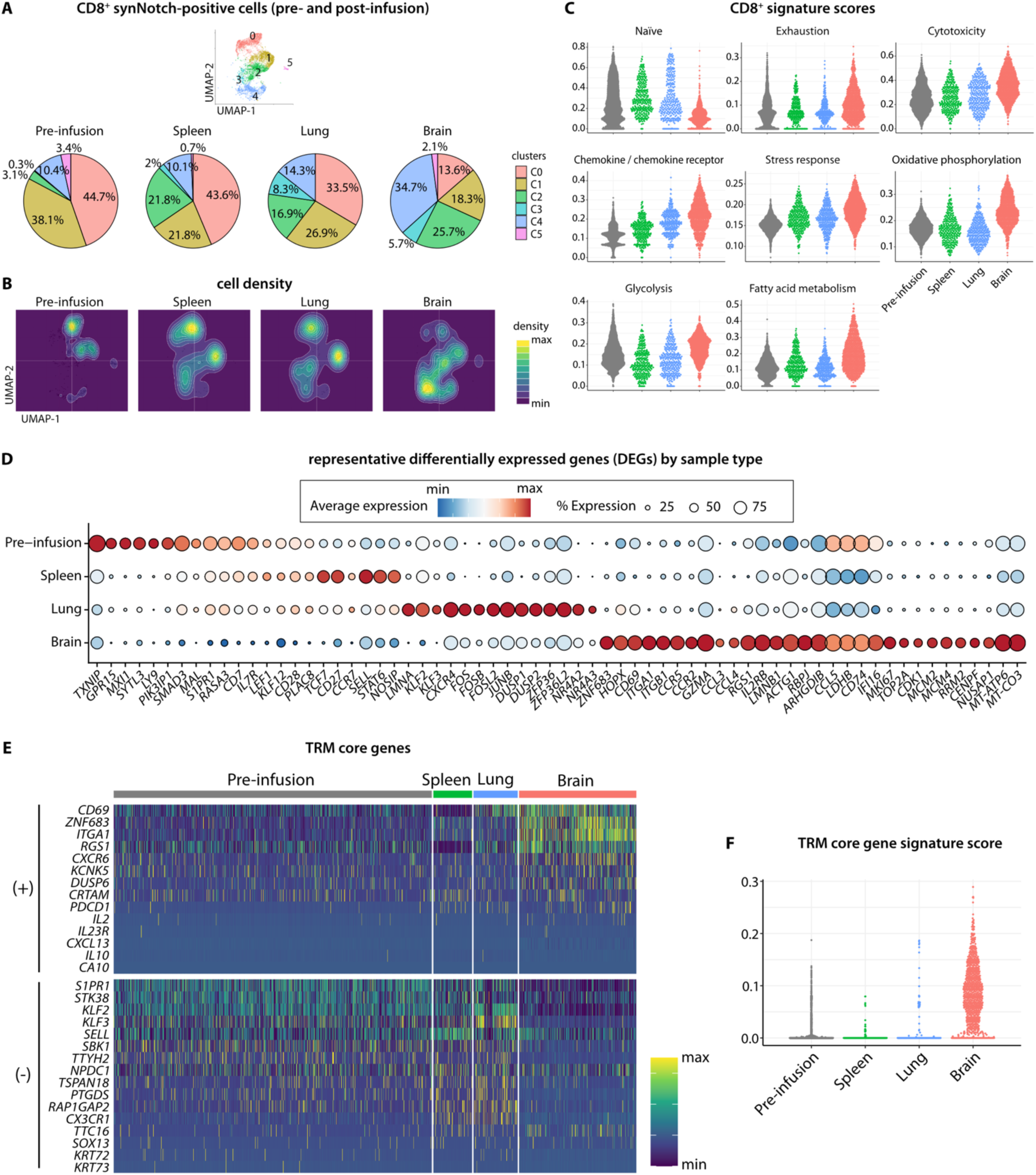
Tracking the transcriptomic evolution of post-infusion synNotch-positive cells. **(A)** Pie charts comparing CD8^+^ cluster proportions across pre-infusion (B-SYNC, unprimed), spleen, lung, and brain samples. **(B)** UMAP density contour plots showing cell distributions for each sample type, with higher-density areas indicated by brighter colors. **(C)** Swarm plots showing representative CD8^+^ T cell signature score distributions across sample types. The full set of 19 analyzed signatures is provided in **Fig. S8**. **(D)** Dot plot showing curated differentially expressed genes (DEGs) across sample types. The complete DEG list is provided in **Supplementary Tables S5–S6**. **(E)** Heatmap showing expression of the tissue-resident memory (TRM) core genes (*37*) in individual cells. **(F)** Swarm plot depicting the distribution of TRM core gene signature scores. Differences among sample types were evaluated using the Kruskal–Wallis test (*P* < 2.5 × 10^-324^).

Differential gene expression analysis further revealed both distinct transcriptomic programs and shared features across tissues (**Fig. 4D; Supplementary Tables S5–S6**). Genes associated with quiescence state and metabolic restraint (*TXNIP, GPR15, MXI1,* and *LY9*) were selectively enriched in the pre-infusion sample. *PIK3IP1*, *SMAD3,* and *MAL* were highly expressed in pre-infusion and moderately expressed in spleen and lung, consistent with naïve or resting phenotypes. Naïve and central-memory-associated genes (*TCF7, LEF1, IL7R, CCR7, CD27, CD28*) were enriched in the spleen, with intermediate expression in pre-infusion and lung samples. Lung-specific features included upregulation of *KLF2, KLF3*, *LMNA,* and *CXCR4*, whereas AP-1 transcription factors (*FOS, FOSB, FOSL2, JUNB*) and early activation genes (*DUSP1/2, ZFP36/ZFP36L2, NR4A2/3*) were most prominently expressed in the lung and present at lower levels in the brain. In contrast, brain-derived cells exhibited a coordinated effector and proliferative program, including mitochondrial genes (*MT-ATP6, MT-CO1/2/3, MT-ND1–5*), cytotoxic markers (*GZMA*, *CCR2/5, CCL3/4/5*), and cell-cycle genes (*MKI67*, *TOP2A*, *CDK1*, *MCM2/4*).

Notably, brain-derived cells displayed upregulation of tissue-residency-associated genes (*CD69, ITGA1, ITGB1, ZNF683/*Hobit, *HOPX*) and reciprocal downregulation of *S1PR1, KLF2/3*, and *MAL* (**Supplementary Tables S5–S6**). In contrast, spleen- and lung-derived cells maintained high *S1PR1* and low *CD69* expression. CD69 is an early marker of tissue residency, whereas S1PR1 (sphingosine-1-phosphate receptor 1) promotes lymphoid egress; their mutually exclusive expression has been well described (*35–37*). TRM core gene signature scoring (*37*) further corroborated these findings, with brain-derived cells exhibiting significantly higher scores (Kruskal–Wallis test, *P* < 2.5 × 10^-324^). These results are consistent with previous observations that CD69 upregulation in infused B-SYNC cells occurs predominantly in the brain (*4*).

Collectively, these findings demonstrate pronounced organ-specific transcriptomic evolution of infused synNotch-CAR-T cells. While spleen- and lung-derived cells largely retain a naïve or quiescent transcriptional profile similar to the pre-infusion state, brain-infiltrating cells undergo coordinated activation characterized by increased cytotoxicity, proliferation, upregulation of metabolic pathways, and acquisition of a TRM-like phenotype. Similar trends were observed in CD4^+^ T cells (**Fig. S9; Supplementary Tables S7–S8**).

### Environmental cues and synNotch-mediated antigen recognition jointly shape transcriptomic remodeling

We next asked whether organ-specific transcriptomic remodeling was driven primarily by environmental cues, synNotch receptor-mediated recognition of the priming antigen, or their combination. To this end, we projected classifier-defined synNotch-negative cells onto the reference UMAP of synNotch-positive cells (**Fig. S10**) and compared synNotch-positive and - negative CD8^+^ cells, including those recovered from the brain. The projection revealed that synNotch-negative cells were largely composed of clusters C0 and C1, corresponding to naïve and quiescent phenotypes. A similar trend was observed in brain-derived cells, although brain-derived cells contained higher fractions of C2–C4 than other tissues.

Organ-specific transcriptomic trends were largely conserved between synNotch-positive and -negative cells in pre-infusion, spleen, and lung samples (**Fig. 5A**), suggesting that remodeling in these tissues was primarily driven by brain-specific environmental cues rather than synNotch-mediated signals. Brain-derived cells, however, exhibited an additional receptor-dependent divergence: synNotch-positive cells moderately upregulated genes associated with activation (*GZMA*, *CCR2/5*, and *CD74*), proliferation (*MKI67*, *TOP2A*, and *CDK1*), and mitochondrial gene expression (*MT-ATP6* and *MT-CO1/3*) compared with synNotch-negative cells. With respect to the TRM core gene signature (*37*), brain-derived synNotch-positive and -negative cells displayed similar profiles, while synNotch-positive cells exhibited slightly higher signature score distributions (Kolmogorov–Smirnov test, *P* = 0.003) (**Fig. S10C**).

**Fig. 5.**
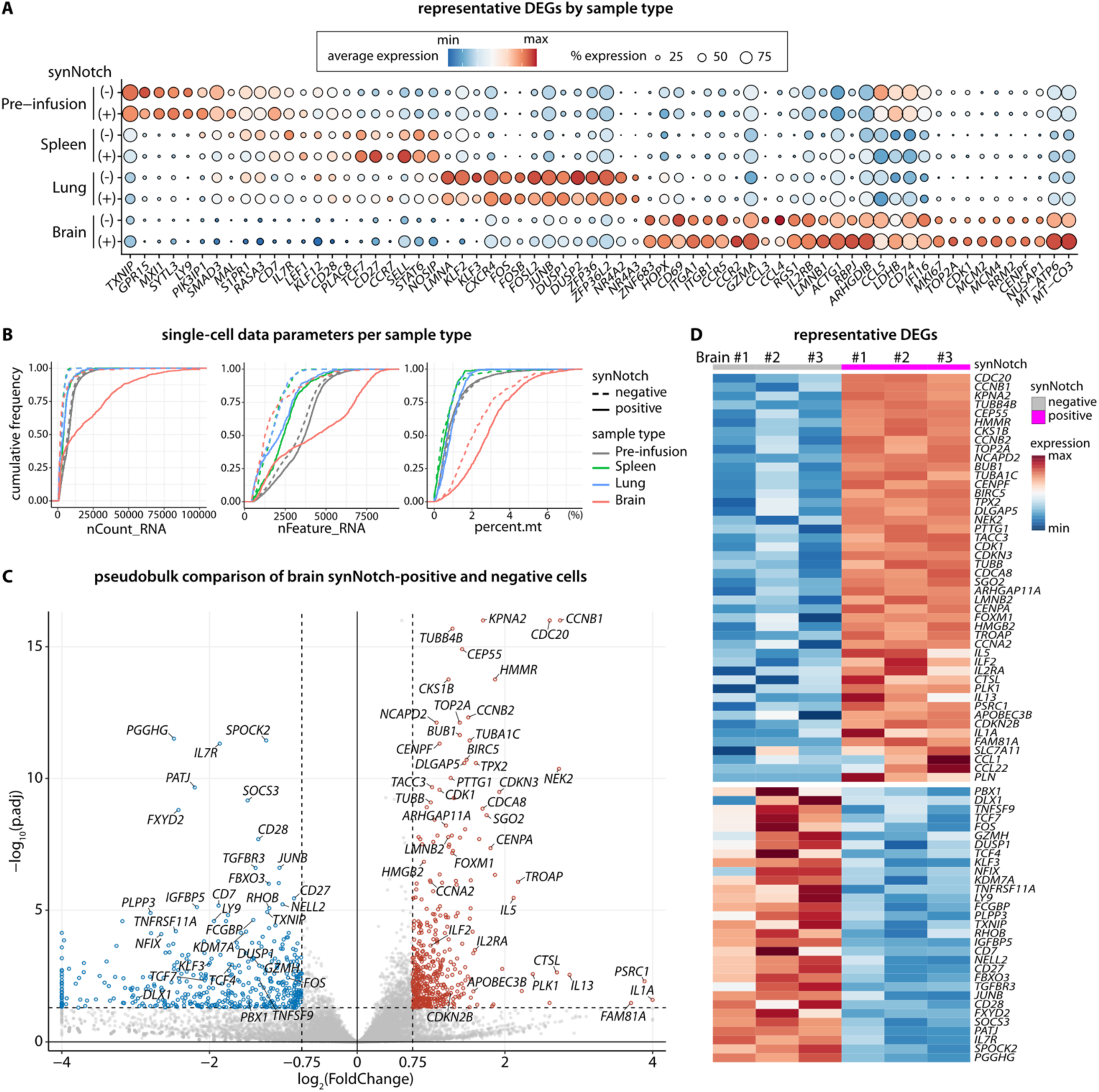
Transcriptomic remodeling by environmental signals and synNotch-mediated antigen recognition. **(A)** Dot plot showing expression of representative differentially expressed genes (DEGs) by sample type (same gene list as in **Fig. 4D)**, stratified by synNotch-positive and -negative cells. **(B)** Empirical cumulative distribution function (ECDF) plots showing the distributions of nCount_RNA (transcript abundance), nFeature_RNA (number of expressed genes), and percent.mt (mitochondrial transcript fraction). Solid and dashed lines indicate synNotch-positive and -negative cells, respectively. **(C, D)** Volcano plot and heatmap summarizing pseudobulk differential expression analysis comparing brain-derived CD8^+^ synNotch-positive and -negative cells. Genes with an adjusted *P* value < 0.05 and |log_2_(Fold Change)| > 0.75 were defined as DEGs. DEGs are highlighted in red or blue, and curated DEGs are labeled in the volcano plot (**C**). Scaled expression levels of these DEGs are shown in the heatmap (**D**). The complete DEG list is provided in **Supplementary Table S9**.

Consistent with this pattern, we observed notable differences in overall transcriptional activity. Brain-derived synNotch-positive cells exhibited substantially higher transcript abundance (nCount_RNA), greater gene expression diversity (nFeature_RNA), and elevated mitochondrial transcript fractions (percent.mt) compared with synNotch-negative cells (**Fig. 5B**). Although similar trends were present in other tissues, the differences were most pronounced in the brain. This aligns with prior observations that overall transcript abundance increases upon T cell activation, as confirmed by reanalysis of an independent dataset comparing resting and activated human T cells (*38*) (**Fig. S11A**) and by differences observed across cell clusters (**Fig. S11B–C**).

A pseudobulk analysis comparing brain-derived synNotch-positive and -negative cells further delineated the molecular basis of this difference. Numerous cell cycle- and proliferation-associated genes (*CDK1, PLK1, CCNB2, CDC20, TOP2A*) were significantly upregulated, whereas naïve and central-memory-associated genes (*IL7R*, *CD7*, *TCF7*, *CD27*, *CD28*) were significantly downregulated in synNotch-positive cells relative to synNotch-negative cells (**Fig. 5C, D; Supplementary Table S9**).

Similar trends were observed in the CD4^+^ population, including environmental influences on cluster composition, transcriptomic profiles, and acquisition of a TRM-like phenotype (**Fig. S10D–F; S12A**). Although differences in transcriptional activity were noted between brain-derived CD4^+^ synNotch-positive and negative cells (**Fig. S12B**), pseudobulk analysis did not reveal significant enrichment of cell-cycle- and proliferation-associated genes, potentially reflecting distinct proliferative profiles of CD8^+^ and CD4^+^ T cells (**Fig. S12C–D**; **Supplementary Table S10**).

Altogether, these findings support a hierarchical model of T cell activation in which environmental cues act as the primary drivers of transcriptomic remodeling, while synNotch-mediated cognate antigen recognition provides an additional amplifying signal, most notably by promoting differentiation and proliferation of CD8^+^ T cells.

### Dynamic regulation and functional implications of CAR transcript expression

A defining feature of the synNotch system is that CAR expression is induced upon successful priming and restricted to sites where the cognate antigen is present. Consistent with this principle, flow cytometric analysis of matched batch samples showed that 51.7% of E-SYNC cells and 40.5% of B-SYNC cells expressed CAR (measured by GFP) following *in vitro* stimulation, whereas only 3.54% and 0.3% of their unprimed counterparts did so, respectively (**Fig. S13A–B**). A central question, therefore, is whether scRNA-seq detection of CAR transcripts reliably reflects priming status.

To address this, we integrated signals across all eight CAR-targeting probes and examined CAR transcript abundance (maxUMI_CAR_) across conditions in synNotch-positive cells. *In vitro*, as expected, primed samples exhibited significantly higher maxUMI_CAR_ values than unprimed samples for both E-SYNC and B-SYNC cells (Kolmogorov–Smirnov test: *D* = 0.37 and 0.22, respectively; *P* < 2.5 × 10^-324^ for both) (**Fig. 6A–C**). *In vivo*, brain-derived cells showed significantly higher maxUMI_CAR_ values for the CAR transcript abundance than spleen- or lung-derived cells (*D* = 0.14, adjusted *P* = 3.2 × 10^-7^ vs. spleen; *D* = 0.20, adjusted *P* = 1.3 × 10^-14^ vs. lung). Notably, low-level CAR transcripts were detectable even in unprimed *in vitro* samples and in spleen and lung samples, indicating baseline transcription independent of priming. Consistent with this observation, classification-based analyses identified substantial fractions of CAR-positive cells in unprimed samples (37.3% in E-SYNC and 31.5% in B-SYNC), compared with 73.9% and 53.0% in primed samples, respectively (**Fig. S14**). Importantly, CAR-detecting probes showed minimal reactivity with UT cells (**Fig. S5**), arguing against cross-reactivity with unrelated transcripts.

**Fig. 6.**
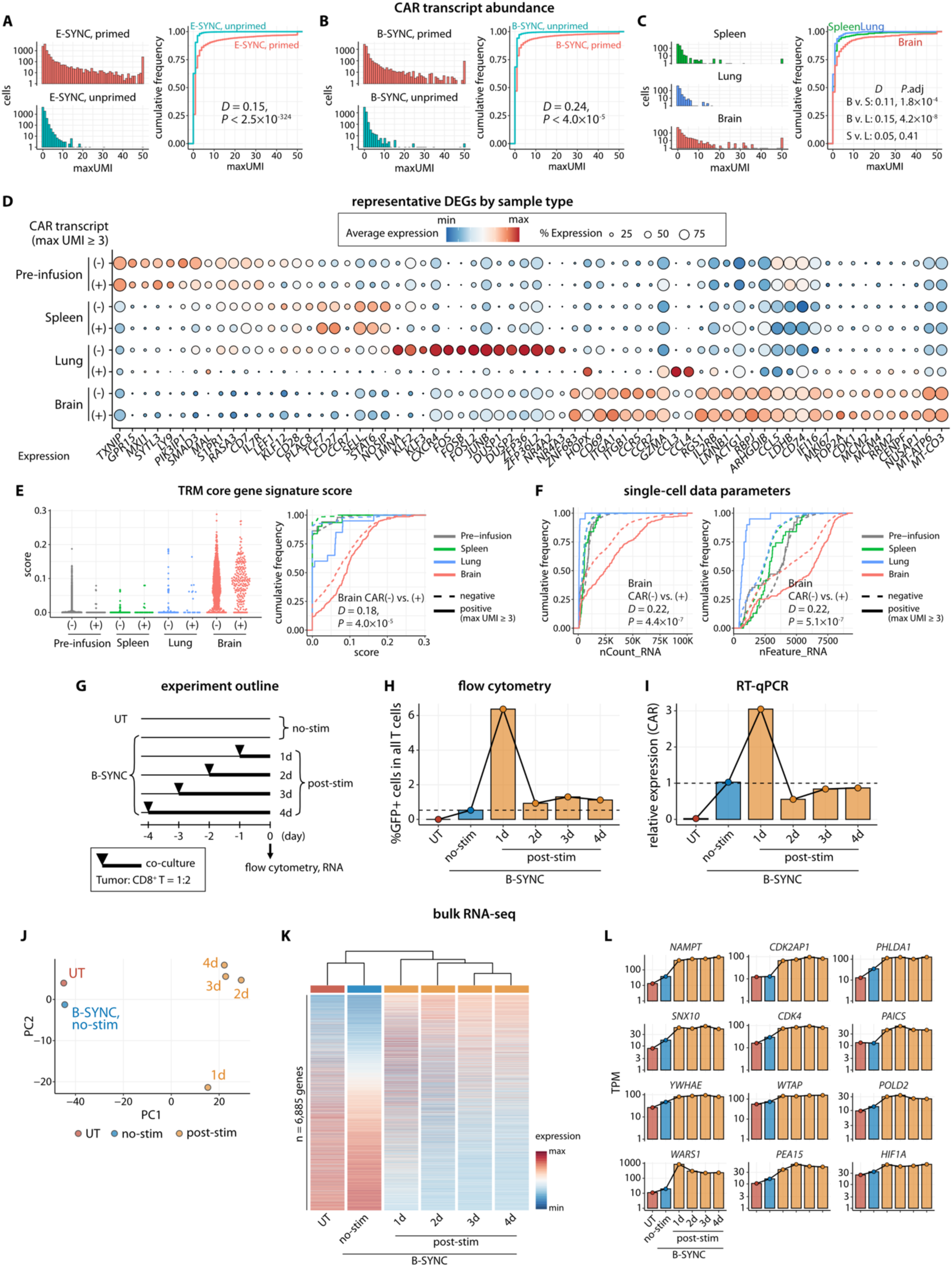
Dynamic regulation and functional implications of CAR transcript expression. (**A–C**) Histograms and empirical cumulative distribution function (ECDF) plots showing distribution of maxUMI_CAR_ values in *in vitro* E-SYNC cells (**A**), *in vitro* B-SYNC cells (**B**), and *in vivo* samples (**C**). In the histograms, the bin width is 1, and the Y-axis is shown on a log_10_ scale. In both histograms and ECDF plots, maxUMI_CAR_ values >50 were capped at 50 for visualization. **(D)** Dot plot showing expression of representative DEGs by sample type (same gene list as in **Fig. 4D and 5A**), stratified by CAR-expressing versus non-expressing groups. **(E)** Swarm plot and ECDF plot depicting the distribution of TRM core gene signature scores. Differences between CAR-positive and -negative cells within the brain were assessed using the Kolmogorov–Smirnov test (*P* = 4.0 × 10^-5^). **(F)** ECDF plots comparing the nCount_RNA and nFeature_RNA values between CAR-expressing and non-expressing cells within the brain-derived CD8^+^ synNotch-positive population. Differences were assessed using the Kolmogorov–Smirnov test. **(G)** Schematic overview of the time-course co-culture experiment of B-SYNC cells with the human GBM cell line GBM6 (BCAN^+^/EphA2^+^/IL13Rɑ2^+^). Co-culture was performed using 24-well plates, with each well containing 0.5 × 10^5^ GBM cells and 1 × 10^6^ CD8^+^ B-SYNC cells. The data presented are representative of two independent experiments using different donors. Additional data are shown in **Fig. S16**. **(H)** Flow cytometry analysis showing the percentage of cells expressing GFP fused to CAR within the live, singlet, non-tumor (mCherry-negative) population. Corresponding scatter plots are shown in **Fig. S16**. **(I)** RT-qPCR data showing CAR transcript abundance across conditions. Relative expression was calculated using the 2^-ΔΔCq^ method, with the B-SYNC no-stimulation (no-stim) condition and the *RPS18* transcript as references. (**J–K**) Bulk RNA-seq principal component analysis (PCA) plot (**J**) and gene expression heatmap (**K**) illustrating transcriptomic similarities and differences across samples. The heatmap displays scaled expression of 6,885 genes expressed in the B-SYNC no-stim sample at transcripts-per-million (TPM) ≥ 10, sorted by the magnitude of expression changes between no-stim and post-stimulation conditions. (**L**) Representative genes minimally expressed without co-culture but upregulated and sustained after co-culture. The full list of gene expression comparisons is provided in **Supplementary Table S13**.

We next evaluated whether threshold-based binary classification could better capture functionally distinct subsets. To explore functional correlates, we applied a stringent threshold (maxUMI_CAR_ ≥ 3) chosen to maximize the difference in CAR-positive frequencies between brain and spleen samples. Under this criterion, 2.6% (118/4,520 cells), 8.5% (60/702 cells), 6.3% (44/699 cells), and 22.2% (262/1,182 cells) of synNotch-positive cells were classified as CAR-high in pre-infusion, spleen, lung, and brain samples, respectively (**Fig. S14**). Within each sample type, CAR-high and CAR-low CD8^+^ cells largely shared similar gene expression profiles. The notable exception was the brain, where CAR-high cells showed upregulation of TRM-associated markers (*CD69*, *ZNF683*, *ITGA1*) and significantly higher TRM core gene signature scores (**Fig. 6D–E**). Brain-derived CAR-high cells also exhibited elevated nCount_RNA and nFeature_RNA values, consistent with increased transcriptional activity (**Fig. 6F**). Similar trends were observed in the CD4^+^ population (**Fig. S15**). Pseudobulk analyses comparing brain-derived CAR-high and CAR-low cells did not identify significant DEGs, with the exception of *CD247* (encoding the CD3ζ chain) (CD8^+^, adjusted *P* = 1.9 × 10^-14^; CD4^+^, adjusted *P* = 9.3 × 10^-15^), consistent with its fusion to the CAR construct (**Supplementary Tables S11–12**). These findings indicate that CAR transcript abundance only partially reflects T cell activation status.

We next examined the kinetics and persistence of CAR transcripts following priming. Our colleagues previously demonstrated that CAR protein expression is transient after synNotch-mediated induction and decays with a half-life of approximately 10 hours, with even faster decay in the presence of cognate antigen, likely due to accelerated CAR endocytosis (*3*). However, the kinetics of CAR transcript expression and their relationship to global cellular transcriptome states have not been thoroughly investigated.

To directly examine transcript kinetics, we performed a time-course co-culture experiment of B-SYNC cells with GBM6 tumor cells (BCAN^+^/EphA2^+^/IL13Rɑ2^+^) (**Fig. 6G**). Because B-SYNC cells rapidly eliminate tumor cells within 24 hours, this system captures the full sequence of events: (i) recognition of the synNotch-specific antigen, (ii) synNotch-driven CAR induction, (iii) recognition of the distinct CAR-specific antigens, (iv) CAR-mediated T cell activation and tumor killing, and (v) a post-eradication phase lacking continued antigen stimulation. Flow cytometry showed that GFP (reporting CAR expression) peaked within 24 h of tumor engagement and declined to near baseline by 48 h (0.53% in no-stim, 6.37% at 24 h, and 0.93% at 48 h; **Fig. 6H and Fig S16**). RT-qPCR confirmed rapid downregulation of CAR transcripts (**Fig. 6I**). Bulk RNA-seq and principal component analysis (PCA) demonstrated that post-stimulation samples clustered distinctly from non-stimulated controls, and that days 2–4 samples formed a coherent cluster separated from day 1 (**Fig. 6J**). Despite declining CAR transcript levels, B-SYNC cells maintained a reprogrammed transcriptional state for at least four days (**Fig. 6K**), including sustained expression of genes involved in metabolic adaptation (*NAMPT*, *WARS1*, *HIF1A, PAICS*), cell-cycle/proliferation program (*CDK4*, *CDK2AP1*, *POLD2*, *WTAP*), and signaling regulators downstream of TCR activation (*PHLDA1, PEA15, YWHAE*) (**Fig. 6L; Supplementary Table S13**). These genes substantially overlapped with transcriptional programs observed in activated CD8^+^ T cells (**Fig. S17A**) and brain-derived synNotch-positive CD8^+^ T cells (**Fig. S17B**).

Collectively, these findings demonstrate that low-level baseline expression of CAR transcripts occurs in the absence of antigen recognition and that CAR transcript abundance is dynamically regulated and partially decoupled from sustained functional activation. Although CAR transcripts are induced upon priming, CAR transcript abundance alone is insufficient to define functional priming. These results underscore the importance of comprehensive transcriptomic profiling to accurately resolve activation and differentiation states in synNotch-CAR-T cells.

## Discussion

We developed a spike-in probe-enhanced scRNA-seq method that addresses a critical unmet need in engineered cell therapy: the ability to simultaneously detect rare CAR-T cells, particularly synNotch-CAR-T cells, and comprehensively profile their functional states at single-cell resolution. By integrating multiple probe signals with ML-assisted classification, our approach achieved robust detection of synNotch-positive cells with high specificity. This strategy allowed us to capture the functional states of engineered T cells across a range of tissues and experimental conditions.

From a technical standpoint, our strategy offers several advantages over poly-A-capture approaches for CAR-T cell tracking (*10–26*). First, the use of multiple spike-in probes targeting the same construct enhances detection robustness without compromising specificity. Second, computational integration of probe-level signals further improves classification accuracy by mitigating technical noise and biological variability. Third, once synNotch-positive cells are determined, standard workflows can be applied for downstream transcriptomic analyses. In the present study, concurrent profiling of host cells, such as myeloid populations, was not possible due to the combination of a xenograft-derived mixed-species dataset and a human transcriptome probe set. In future clinical trial samples, however, this limitation will not apply, as all cells will be human-derived, and simultaneous profiling of both infused cells and host immune populations should therefore be fully achievable. This probe-based approach has inherent limitations; it cannot be combined with TCR characterization (V[D]J sequencing), and the absence of raw read information precludes RNA-processing-related analyses, such as RNA velocity (*39*). Nonetheless, the probe-based method offers several practical benefits, including efficient multiplexing and reduced per-sample cost, as well as increased flexibility in sample handling timelines enabled by fixation and cryopreservation. These features are particularly advantageous for analyzing longitudinal samples from multiple patients.

The efficacy and tissue specificity of synNotch-CAR-T cells have been previously characterized in our previous study, primarily at the protein level (*4*). Our single-cell analyses orthogonally validated these findings, revealing substantial transcriptional remodeling of synNotch-CAR-T cells after *in vivo* infusion, with distinct patterns emerging across organs.

Brain-derived cells, regardless of synNotch status as of the time of harvesting, displayed pronounced activation signatures characterized by elevated expression of effector molecules, chemokines, exhaustion markers, metabolic genes, and tissue-resident memory (TRM) markers. In contrast, spleen-derived cells retained a naïve-like or quiescent phenotype, whereas lung-derived cells exhibited prominent AP-1-related transcriptional activity (e.g., upregulation of FOS/JUN family genes), potentially reflecting early priming events prior to brain infiltration (*40*). Interestingly, beyond environmental influences, synNotch-mediated cognate antigen recognition also contributed: synNotch-positive cells within the brain exhibited increased expression of cell-cycle markers and reduced expression of naïve markers compared to synNotch-negative cells in the brain. Together, these data support a hierarchical model in which tissue-dependent cues establish the primary differentiation landscape, while synNotch-mediated antigen recognition provides an additional proliferative and activation boost.

In particular, strong induction of a TRM phenotype was observed in brain-infiltrating T cells, exemplified by upregulation of *CD69* and coordinated downregulation of *S1PR1*, with this trend more pronounced in synNotch-positive cells. This observation is notable, given that synNotch-CAR-T cells are designed to reside in the brain and that TRM cells are associated with improved prognosis in multiple cancer types (*34, 41, 42*). It is worth noting that expression of *ITGAE*/CD103, a canonical marker associated with TRM cells, could not be assessed, as it was not included in the premanufactured whole-transcriptome probe set. In future studies, the incorporation of custom spike-in probes targeting *ITGAE* would enable a more comprehensive assessment of TRM maturation dynamics.

Beyond tissue residency, an important translational consideration is how to interpret CAR expression in clinical samples. Because low-level baseline CAR transcripts are detectable even in the absence of antigen recognition and are dynamically regulated upon priming, CAR transcript abundance alone may not directly reflect the functional activation status of synNotch-CAR-T cells. Detection of CAR transcripts in tissues outside the brain was therefore not unexpected, as the E-/B-SYNC circuits were tuned such that baseline transcript levels do not reach functionally significant CAR expression capable of inducing on-target, off-tumor toxicity. Nevertheless, binary classification into CAR-expressing versus non-expressing cells did not align with global scRNA-seq transcriptomic profiles, which demonstrated that many CAR transcript-negative cells in the brain exhibited activated and functional states. It is plausible that these cells had previously expressed CAR transcripts but subsequently downregulated them, with only a subset retaining detectable transcript levels at the time of analysis, as partially recapitulated in our time-course experiments. Fundamental questions remain regarding the relationship between transcript and protein expression, functional activation thresholds, promoter dynamics, and their temporal regulation, underscoring the complexity of interpreting CAR transcript abundance at a single time point. Accordingly, comprehensive transcriptomic profiling is likely to be more informative than reliance on CAR expression alone for identifying successfully primed, functional synNotch-CAR-T cells *in vivo*.

We acknowledge several limitations in this study. First, the number and quality of recovered cells varied across samples. *In vivo* tissues yielded fewer viable cells and contained mouse-derived cells that had to be computationally filtered, which affected the data balance.

Future experimental designs should account for this variability. Second, all data were derived from preclinical models; thus, validation using clinical specimens will be essential. While we observed tissue-dependent remodeling of B-SYNC cells, the xenograft setting imposes additional constraints, as human tumor-derived proteins are present only in tumor-bearing brain tissue. Third, synNotch gene silencing data were unavailable for the corresponding cell batches. Although the rate of gene silencing has been empirically estimated at approximately 20–40%, the lack of precise, matched data precluded accurate sensitivity calculations for synNotch detection models. Fourth, synNotch gene silencing following infusion is theoretically possible but technically challenging to evaluate and has not been systematically assessed.

Consequently, a key unsolved question is whether synNotch-negative cells detected in the brain represent false-negative cells (i.e., cells not detected by scRNA-seq) or cells that underwent gene silencing either before infusion, during circulation, or after migration into the brain. Addressing these possibilities will require further well-designed investigations.

In summary, our spike-in probe-enhanced scRNA-seq approach provides a robust and scalable platform for accurate detection and high-resolution transcriptomic characterization of synNotch-CAR-T cells *in vivo*. By resolving functional adaptations and differentiation trajectories after infusion, this strategy establishes a generalizable framework for tracking engineered immune cells within complex tissue environments. Beyond synNotch-CAR-T cell applications, the method holds broad potential for clinical monitoring and mechanistic studies of engineered cell therapies in which defined transgene sequences can be leveraged for detection using spike-in probes.

## Materials and Methods

### Study design

The objective of this study was to develop a method for detecting and characterizing synNotch-CAR-T cells after infusion using the 10x Genomics FLEX (probe-based) platform. Custom spike-in probes (n = 20) were designed to target synNotch transcripts specific to E-SYNC and B-SYNC cells, as well as the shared CAR transcript. The approach was evaluated using scRNA-seq across 16 preclinical samples (5 *in vitro* and 11 *in vivo*).

SynNotch-positive cells were identified using spike-in probe signals combined with machine-learning (ML)-assisted classification. Downstream transcriptomic analyses were performed to assess cell state, differentiation trajectories, tissue-specific differences, and synNotch receptor-mediated signaling. The relationship between CAR transcript abundance and transcriptional state was also evaluated.

### Construct design and generation

SynNotch receptors were built by fusing the various scFv sequences (Sidhu lab or patents) to mouse Notch1 (NM_008714) minimal regulatory region (residues 1427 to 1752) and Gal4 DBD VP64 (*3, 4*). All synNotch receptors contain N-terminal CD8a signal peptide (MALPVTALLLPLALLL HAARP) for membrane targeting and α-myc-tag (EQKLISEEDL) for detecting surface expression with α-myc A647 (Cell Signaling Technology, catalog no. 2233). See Morsut *et al* (*5*) for the synNotch sequence. Receptors were cloned into a modified pHR’SIN:CSW vector containing a PGK or SFFV promoter. The pHR’SIN:CSW vector was also used to make response element plasmids with five copies of the Gal4 DNA-binding domain target sequence (GGAGCACTGTCCTCCGAACG) upstream from a minimal CMV promoter. Response element plasmids also contain a PGK promoter that constitutively drives BFP expression to easily identify transduced T cells. CARs were built by fusing IL-13 Mutein [E13K,K105R]-G4Sx4-EphA2 scFv (*3*) to the hinge region of the human CD8α chain and transmembrane and cytoplasmic regions of the human 4-1BB, and CD3ζ signaling domains. Inducible CAR constructs were cloned into a site 3’ to the Gal4 response elements and minimal CMV promoter. CARs were tagged c-terminally with GFP or N-terminally with myc tag to verify surface expression.

### Primary human T cell isolation and culture

Primary CD4^+^ and CD8^+^ T cells were isolated from donor blood after apheresis by negative selection (STEMCELL Technologies). Blood was obtained from StemExpress or Allcells, as approved by the University of California San Francisco (UCSF) institutional review board. T cells were cryopreserved either in Cellbanker 1 or in RPMI-1640 with 20% human AB serum (Valley Biomedical) and 10% dimethyl sulfoxide. After thawing, T cells were cultured in human T cell medium consisting of X-VIVO 15 (Lonza), 5% human AB serum, 55 µM β-mercaptoethanol, and 10 mM neutralized *N*-acetyl-l-cysteine supplemented with 30 units/ml IL-2.

### Lentiviral transduction of human T cells

Pantropic vesicular stomatitis virus G (VSV-G) pseudotyped lentivirus was produced through transfection of Lenti-X 293T cells (Takara Bio, catalog no. 632180) with a pHRʹSIN:CSW transgene expression vector and the viral packaging plasmids pCMV and pMD2.G using Fugene HD (Promega) or TransIT-VirusGen (Mirus Bio). Primary T cells were thawed the same day and, after 24 hours in culture, were stimulated with 25 µl of anti-CD3/CD28 coated beads (Dynabeads Human T-Activator CD3/CD28, Gibco) per 1 × 10^6^ T cells. In some instances, freshly isolated T cells were used. At 48 hours, viral supernatant was harvested. Primary T cells were exposed to the lentivirus for 24 hours or spinoculated on retronectin-coated plates. At day 5 after T cell stimulation, Dynabeads were removed and T cells were sorted with a BD Biosciences FACSARIA Fusion or Sony SH800S Cell Sorter. T cells exhibiting basal CAR protein expression (measured by GFP) were gated out during sorting. T cells were expanded until rested and could be used in assays.

### Design of spike-in probes

Spike-in probes were designed to target three regions within the synNotch-CAR vectors: (1) the E-SYNC synNotch receptor targeting EGFRvIII (1,692 bp); (2) the B-SYNC synNotch receptor targeting BCAN (1,698 bp); and (3) the anti-EphA2/IL13ɑ2 dual CAR shared by both E-SYNC and B-SYNC (1,173 bp) (**Fig. 1**). Following the manufacturer’s instructions (*CG000621_CustomProbeDesign_RevD,* 10x Genomics) (*29*), all eligible 50-base probe sequences were identified (n = 574 in total). A BLAST search was then performed for all candidate 25-base left- and right-hand side (LHS or RHS) sequences using the following parameters: database: *core_nt* (core nucleotide BLAST database); organism: *Homo sapiens* (taxid: 9606); program: *megablast* (highly similar sequences). Candidate LHS and RHS sequence pairs were ranked based on match score and alignment length. To maximize transcript coverage and prevent overlap among probe target regions, candidates were manually curated and prioritized. A subset of 45 probe candidates was selected for specificity testing using reverse transcription quantitative PCR (RT-qPCR), as described below. Based on RT-qPCR results and probe distribution across target transcripts, a final set of 20 probes was selected (**Supplementary Table S1**).

### Reverse transcription-quantitative PCR (RT-qPCR)

Total RNA was extracted using the RNeasy Mini Kit (Qiagen, 74106) with on-column DNase treatment (RNase-Free DNase Set, Qiagen, 79254) according to the manufacturer’s instructions. Complementary DNA (cDNA) was synthesized using qScript Ultra SuperMix (Quantabio, 95217-100). To evaluate probe candidates, corresponding pairs of 25-base primers were designed, and *GAPDH* was used as a reference gene (Forward: 5’-CATCACTGCCACCCAGAAGACTG-3’; Reverse: 5’-ATGCCAGTGAGCTTCCCGTTCAG-3’).

For the B-SYNC cell time-course experiment, the following primers were used: CAR forward: 5’-TGACTGCGGGCATGTATTGT-3’; CAR reverse: 5’-CCCTACTTGAGGCGCCATAG-3’; RPS18 forward: 5’-GCAGAATCCACGCCAGTACAAG-3’; RPS18 reverse: 5’-GCTTGTTGTCCAGACCATTGGC-3’. RT-qPCR reactions (10 µl) contained 5 µl of PerfeCTa SYBR Green FastMix ROX (Quantabio, 95073-012), 0.5 µl each of forward and reverse primers, 2 µl of RNase-free water, and 2 µl of normalized cDNA input. Reactions were run in triplicate on a MicroAmp™ Optical 384-well reaction plate (Applied Biosystems, 4309849) using a QuantStudio 5 instrument (Applied Biosystems). Quantification cycle (C_q_) values and amplification curve data were exported using the QuantStudio Design and Analysis v2 software. Only reactions labeled as ‘Amp’ (*Target amplified*) were included in downstream analysis. Relative expression levels were calculated using the 2^-ΔΔCq^ method, normalized to *GAPDH* or *RPS18* and to control samples as specified in the corresponding figure legends.

### Spike-in probes

For each of the 20 selected probe targets, LHS and RHS probe sequences were determined according to the manufacturer’s specifications (*29*) and synthesized as 10 pmol oPools oligos for each of 16 barcodes (BC001–BC016) by Integrated DNA Technologies (IDT). Detailed probe information is provided in **Supplementary Table S1**.

### Cell lines

GBM6 PDX cells (gift from Frank Furnari, Ludwig Institute and UCSD) were lentivirally transduced to stably express mCherry, and enhanced firefly luciferase (fluc) under control of the spleen focus-forming virus (SFFV) promoter and sorted as previously shown (*3*). GBM6 cells were cultured in DMEM/F12 medium, with supplements of B27, epidermal growth factor (EGF, 20 µg/ml), fibroblast growth factors (FGF, 20 µg/ml), and heparin (5 µg/ml).

### *In vitro* stimulation of synNotch-CAR-T cells for scRNA-seq analysis

To prepare the E-SYNC primed sample, 5 × 10^4^ cells were co-cultured for 24 hours with EGFRvIII-coated beads (ACROBiosystems, MBS-K020). To prepare the B-SYNC primed sample, 2 × 10^4^ cells were cultured for 24 hours on wells coated with Laminin (100 µg/ml, Thermo Fisher Scientific,23017015) and recombinant BCAN (25 µg/ml; R&D Systems, 7188-BC-050 and 4009-BC-050). Cells were analyzed by flow cytometry using an Attune NxT (Thermo Fisher Scientific), and data processed with FlowJo v10.8.1 (TreeStar).

### Time-course co-culture of GBM6 cells and B-SYNC cells

B-SYNC CD8^+^ cells were prepared as described above. For co-culture experiments, 5 × 10^4^ GBM6 cells were seeded on day -5 (relative to the day of sample collection, day 0) in wells coated with Laminin in a 24-well plate. On days -4, -3, -2, and -1, 5 × 10^4^ CD8^+^ B-SYNC cells were added to designated wells at a 1:1 effector-to-target (E:T) ratio and maintained until sample collection. This staggered co-culture design enabled reconstruction of temporal transcriptional dynamics following synNotch activation. At the time of collection, a fraction of cells was analyzed by flow cytometry, and the remaining cells were subjected to RNA extraction. RNA was analyzed by RT-qPCR and bulk RNA-seq. In a repeated experiment (**Fig. S16B–D**), 5 × 10^4^ GBM6 cells were seeded on day -4, and co-cultured with 1 × 10^5^ CD8^+^ B-SYNC cells (2:1 E:T ratio), with co-culture initiated on days -3, -2, -1, and 12 hours before collection.

### *In vivo* mouse experiments

All mouse experiments were conducted in accordance with Institutional Animal Care and Use Committee (IACUC)-approved protocols. For xenograft experiments, 2 × 10^4^ GBM6-luc-mCherry cells were dissociated into single-cell suspensions and inoculated intracranially into 6- to 8-week-old female NCG mice (Charles River Laboratories). Following anesthesia with 1.5% isoflurane, stereotactic tumor cell inoculation (injection volume: 2 µl) was performed with the injection site 2 mm right to the bregma, and 3 mm below the brain surface. Mice were treated with analgesics before surgery and for 3 days postoperatively and monitored for adverse symptoms in accordance with IACUC guidelines. Tumor engraftment and progression were evaluated by bioluminescence imaging (BLI) using an IVIS Xenogen 2000 following intraperitoneal injection of D-luciferin (1.5 mg in 100 µl; GoldBio). Upon confirmation of tumor establishment (average radiance > 3.5 × 10^3^ photons/s/cm²), mice were treated with either B-SYNC or UT cells (3 × 10^6^ CD8^+^ cells and 3 × 10^6^ CD4^+^ cells) via tail vein injection in 100 µl PBS, as indicated in **Fig. 2A**.

### Isolation of mononuclear cells from *in vivo* tissues

On day 20 after tumor inoculation (day 11 after T cell infusion), mice were euthanized and transcardially perfused with cold PBS, and the spleen, lung, and brain were harvested. Brains were mechanically minced and dissociated on a gentleMACS™ Octo Dissociator with Heaters (Miltenyi 130-096-427) using the brain tumor dissociation program (‘37C_BTDK_1’) in the presence of a digestion mix containing Collagenase D (30 mg/ml; Sigma, C5138), DNase I (10 mg/ml; Sigma, DN25), and soybean trypsin inhibitor (20 mg/ml). The resulting homogenate was filtered through a 70 µm cell strainer and resuspended in 30% Percoll (GE Healthcare), underlaid with 70% Percoll and centrifuged for 30 min at 650 × g. Brain-infiltrating mononuclear cells were recovered from the 30%–70% interface. Spleens were dissociated into single-cell suspensions, and red blood cells were lysed with ACK lysis buffer (Lonza, 10-548E) according to the manufacturer’s instructions. Lung lobes were digested on a shaker in RPMI media containing 10% FBS, collagenase IV and DNase I for 30 min at 37 °C and filtered through a 70 µm cell strainer. The resulting homogenate was resuspended in 40% Percoll, carefully layered on top of 80% Percoll and centrifuged for 20 min at 800 × g. The mononuclear cells were collected from 40%–80% interface.

### Fixation and cryopreservation of cells

Cell fixation was performed using the Chromium Next GEM Single Cell Fixed RNA Sample Preparation Kit (10x Genomics, PN-1000414) according to the manufacturer’s instructions (*43*). For both *in vitro* and *in vivo* samples, cells were washed twice with cold PBS, resuspended in 4% formaldehyde diluted in the provided Fix & Perm Buffer, and incubated overnight at 4°C. After 16–24 hours, the fixation reaction was quenched with the supplied Quenching buffer. After cell counting, the supplied Enhancer (10% of the total volume) and 50% glycerol (final concentration of 10%; Invitrogen, 15514-011) were added, and samples were cryopreserved at -80°C until library preparation (stored for up to 2 months in this study).

### Library preparation and sequencing

Single-cell library preparation was performed by the UCSF Functional Genomics Core Facility according to the manufacturer’s instructions (*44*). For probe hybridization, 40 nM total (LHS and RHS combined) of each custom spike-in probe was mixed with 20 µl Human WTA Probes bearing the corresponding barcode (10x Genomics, PN-2000495–2000510). After hybridization, all 16 samples were pooled and loaded onto a 10x Chromium Next GEM chip and partitioned into gel bead-in-emulsion (GEM) microdroplets. The experiment targeted 128,000 total cells (∼8,000 cells per sample). Paired-end 100-bp reads were generated on an Illumina NovaSeq X using a 25B flow cell at the UCSF Center for Advanced Technology (CAT). FASTQ files were processed with *cellranger multi* (Cell Ranger v9.0.1) using the Chromium_Human_Transcriptome_Probe_Set_v1.1.0_GRCh38-2024-A probe set supplemented with custom spike-in probe definitions.

## Single-cell RNA-seq data analysis

### Detection of human cells

The filtered count matrix generated by Cell Ranger was processed using Seurat (v4.0.3) in R (v4.1.3) (*45*). A total of 73,193 cells were initially detected across 16 samples; After quality control (QC) filtering (nFeature_RNA ≥ 400, 500 ≤ nCount_RNA ≤ 200,000, percent.mt ≤ 7.5), 71,541 cells were retained. Unique molecular identifier (UMI) counts were normalized and scaled using SCTransform (v0.3.5). Dimensionality reduction (RunPCA, RunUMAP), neighbor graph construction (FindNeighbors), and clustering (FindClusters, resolution = 0.1) identified 10 clusters (C0–C9). Based on QC metrics and expression of human T cell markers (**Fig. 2D– F**), four clusters (C0–C2 and C7) were classified as human cell clusters. The brain-UT sample (containing only two human cells after filtering) was excluded from downstream analyses.

Heterotypic doublets were identified and removed using scDblFinder (*46*), and only singlet cells passing QC were included in subsequent analyses.

### Classification of synNotch-positive and -negative cells

Three spike-in probes (P09, P10, P12) were excluded due to poor performance (**Fig. S2**). Raw UMI counts were used as input without any normalization or scaling. At the single-probe level, cells were considered positive if they had ≥1 UMI for that probe.

For multi-probe classification of synNotch-positive versus -negative cells, 13 probes (P01–P04, P11, P13–P20) were used to distinguish *in vitro* E-SYNC cells (both unprimed and primed) from UT, whereas 16 probes (P01–P08, P13–P20) were used to distinguish *in vitro* B- SYNC cells (both unprimed and primed) from UT and to analyze *in vivo* samples. The following classifiers were used:

(1) **Maximum UMI counts (maxUMIsynNotch):** the maximum UMI count across selected probes was calculated for each cell, and distributions of maxUMI_synNotch_ values were compared between E-SYNC and UT or between B-SYNC and UT.
(2) **Machine-learning (ML)-based classifiers:** using the R package *caret*, four algorithms were applied: (i) generalized linear model (GLM; logistic regression), (ii) elastic net-regularized logistic regression (GLMnet), (iii) random forest (RF), and (iv) support vector machine (SVM). E-SYNC or B-SYNC cells were labeled as positive and UT cells as negative. Model performance was assessed using repeated k-fold cross-validation (10 folds, 10 repeats), with class probabilities estimated and performance summarized by the area under the receiver operating characteristic curve (AUC) via the twoClassSummary function. Predictions from all resampling iterations were retained for downstream analysis. SynNotch-positive probabilities were computed for each cell.

Thresholds for each approach (maxUMI_synNotch_ values or probabilities) were determined using Youden’s J statistic to balance sensitivity and specificity (*30*). Cells were defined as synNotch-positive if all approaches concordantly classified them as positive; all remaining cells were defined as synNotch-negative (**Fig. S4**). Across models, classification outputs were highly concordant, yielding consistent cell labels across methods.

### Single-cell transcriptomic profiling analyses

SynNotch-positive (n = 29,465 cells) and synNotch-negative (n = 13,222 cells) populations from 13 non-UT samples (4 *in vitro* and 9 *in vivo*) were analyzed. CD4^+^ and CD8^+^ T cell populations were defined based on normalized expression of *CD4*, *CD8A*, and *CD8B2*. Cells were classified as CD4^+^ if *CD4* expression exceeded *CD8A*, as CD8^+^ if either *CD8A* or *CD8B2* exceeded *CD4*, and as undetermined if neither condition was met.

Each synNotch-positive CD8^+^ and CD4^+^ dataset was divided into five subsets: (1) E-SYNC; (2) B-SYNC (*in vitro*); (3) spleen; (4) lung; (5) brain (*in vivo* B-SYNC). Each subset was normalized and scaled using SCTransform and projected into a lower-dimensional space using RunPCA in Seurat. The five subsets were then integrated using the SCT integration workflow with default parameters (*47*). The integrated data were used for dimensionality reduction (RunPCA, RunUMAP) and neighbor graph construction (FindNeighbors) in Seurat. Louvain clustering based on a shared nearest-neighbor graph was performed using FindClusters (resolution = 0.15 for CD8^+^ and 0.18 for CD4^+^), identifying six clusters in CD8^+^ (C0–C5) and four clusters in CD4^+^ (C0–C3). Pseudotime analysis was performed using slingshot (v2.7.0) on CD8^+^ clusters C0–C4 and the full CD4^+^ dataset. Differential gene expression analysis was performed using FindAllMarkers on the SCT data slot to characterize transcriptomic programs across clusters. Results are provided in **Supplementary Tables S3–S4**, and curated gene expression data are shown in **Fig. 3G and Fig. S7C**. Gene signature analysis was performed on normalized values in the RNA data slot using the ScoreSignature_UCell function in UCell (v1.3.1) (*48*), with 19 CD8^+^ and 17 CD4^+^ human tumor-infiltrating T cell gene signatures (*49*). Projection of cells onto reference single-cell atlases was performed using the Run.ProjecTILs function in ProjecTILs (v3.1.3), with reference atlases of human CD8^+^ T cells (*32, 33*) (available at: https://doi.org/10.6084/m9.figshare.23608308) and CD4^+^ T cells (*32, 34*) (available at: https://doi.org/10.6084/m9.figshare.21981536.v1), respectively.

### Comparison of transcriptomic profiles across sample types

To investigate post-infusion transcriptomic remodeling of synNotch-positive cells, the *in vitro* B-SYNC unprimed sample (pre-infusion) and *in vivo* samples (spleen, lung, and brain) were analyzed (CD8^+^, n = 4,096; CD4^+^, n = 2,326). Cluster identities were retained from Louvain clustering and visualized in 2D UMAP space. Differential gene expression analyses were performed on normalized gene expression data (RNA assay slot) using the FindAllMarkers function in Seurat. Two comparisons were conducted: (1) pre-infusion vs. brain and (2) spleen vs. lung vs. brain. Results are provided in **Supplementary Tables S5–S8**, and curated gene expression profiles are shown in **Fig. 4D** and **Fig. S9D**.

Expression of the reported tissue-resident memory (TRM) core genes (*37*) was assessed across samples. Among the reported set of 31 genes, *ITGAE*, *D4S234E*, and *FAM65B* were not detected in the present dataset and were therefore excluded from analysis. The remaining 28 genes were used to compute TRM gene signature scores using the ScoreSignature_UCell function in UCell. Among these genes, 12 (*CA10, ITGA1, IL2, IL10, CXCR6, CXCL13, KCNK5, RGS1, CRTAM, DUSP6, PDCD1,* and *IL23R*) were defined as positive contributors to the TRM signature, whereas the remaining 16 genes (*STK38, TTC16, SELL, KLF3, KLF2, SBK1, TTYH2, NPDC1, KRT72, S1PR1, SOX13, KRT73, TSPAN18, PTGDS, RAP1GAP2,* and *CX3CR1*) were defined as negative contributors.

### Comparison between synNotch-positive and -negative cells

CD8^+^ and CD4^+^ synNotch-negative cells were processed with SCTransform, RunPCA, RunUMAP, and FindNeighbors in Seurat and then projected onto the corresponding synNotch-positive reference datasets using the Run.ProjecTILs function in ProjecTILs. SynNotch-positive and -negative cells from pre-infusion, spleen, lung, and brain samples were merged within CD8^+^ and CD4^+^ populations, respectively, and subsequently normalized and scaled using the RNA assay slot.

For CD8^+^ and CD4^+^ populations, pseudobulk analysis was performed to compare brain-derived synNotch-positive and -negative cells. UMI counts were aggregated per sample (mouse 1–3) and by synNotch-positive and -negative populations. After filtering lowly expressed genes, differential gene expression analysis was conducted using DESeq2 (v1.32.0). Results are provided in **Supplementary Tables S9–S10**. Genes with adjusted *P* values < 0.05 and |log_2_(fold change)| > 0.75 were defined as significant, of which curated ones were labeled in volcano plots and displayed in heatmaps (**Fig. 6C–D** and **Fig. S12C–D**).

### Assessment of CAR transcript expression

To assess CAR transcript expression, 8 probes (P13–P20) were used. The following approaches were applied:

(1) **Maximum UMI counts (maxUMICAR):** the maximum UMI count across selected probes was calculated for each cell, and distributions of maxUMI_CAR_ values were compared across conditions, including E-SYNC primed vs. unprimed, B-SYNC primed vs. unprimed, and among spleen, lung, and brain samples.
(2) **Machine-learning (ML)-based classifiers:** ML classifiers were implemented as described for synNotch detection. Four algorithms were applied, including (i) GLM, (ii) GLMnet, (iii) RF, and (iv) SVM. E-SYNC and B-SYNC primed cells were labeled as positive and unprimed cells as negative. Model performance was assessed using repeated k-fold cross-validation (10 folds, 10 repeats), with class probabilities estimated and performance summarized by the AUC via the twoClassSummary function. Predictions from all resampling iterations were retained for downstream analysis. Probabilities of CAR positivity were computed for each cell.

Additionally, based on the maxUMI_CAR_ approach, different cutoff values were evaluated, and a threshold of maxUMI_CAR_ ≥ 3, which maximized the difference in CAR-positive frequencies between spleen and brain samples, was selected for threshold-based binary classification (CAR-high vs. CAR-low). Pseudobulk analysis was performed to compare brain-derived CAR-high and CAR-low cells. UMI count data were aggregated per sample (mouse 1– 3) and by CAR-high and CAR-low populations. After filtering lowly expressed genes, differential gene expression analysis was conducted using DESeq2. Results are provided in **Supplementary Tables S11–S12**.

### Bulk RNA-seq data acquisition and analysis

RNA integrity was assessed using an Agilent Bioanalyzer 5400 and confirmed to have a RIN (RNA Integrity Number) value of 7 or greater (range: 7–9.4). RNA quality assessment, library preparation, and sequencing were performed by Novogene (Sacramento, CA). Briefly, mRNA was purified from total RNA using poly-T oligo-attached magnetic beads. After fragmentation, first-strand cDNA synthesis was performed using random hexamer primers, followed by second-strand cDNA synthesis. Libraries were constructed through end repair, A-tailing, adapter ligation, size selection, amplification, and purification. Quality control was conducted using Qubit and real-time PCR for quantification, and Bioanalyzer for size distribution assessment. Finally, quantified libraries were pooled and sequenced on an Illumina NovaSeq X Plus PE150 platform at Novogene, based on effective library concentration and desired sequencing depth.

### Bulk RNA-sequencing data processing and analysis

Quality checking, trimming, and barcode removal were performed using fastp (v0.20.0) with default parameters. FASTQ reads were aligned to the human reference genome GRCh38.p14 using STAR (v2.7.9a), with transcriptome annotation provided by gencode.v49.annotation.gtf. Sorting and indexing were performed using samtools (v1.22.1), and gene-level expression counts and transcripts per million (TPM) values were quantified using StringTie (v2.0). All downstream analyses were conducted using R (v4.1.3). Differential expression (DE) analysis was performed to compare the B-SYNC, no-stimulation sample (n = 1) with co-cultured samples (n = 4) using the DESeq2 R package (v1.32.0). Statistical significance was not formally assessed due to the absence of biological replicates.

## Data visualization

Graphical illustrations were generated using the BioRender web tool. Data visualization was performed using R (v4.1.3) with the ggplot2 package (v3.3.6). All figures were rendered using Affinity Designer (Serif, v1.10.5).

## Statistical analysis

Statistical analysis was performed using R version 4.1.3 as described in the figures and legends.

## Data availability

Transcriptome datasets generated during this study will be deposited in the NCBI Gene Expression Omnibus (GEO) and made publicly available upon publication.

## Code availability

The code used for data analysis will be made publicly available on GitHub upon publication in a peer-reviewed journal.

## Acknowledgments

We thank the following individuals and organizations: the study participants and their families; Samantha Shelton and Brian Chan of 10X Genomics for technical assistance; Lenka Maliskova, Armita Norouzi, and Walter Eckalbar of UCSF Genomics CoLab for the single-cell RNA-sequencing library preparation; Sequencing was performed at the UCSF Center for Advanced Technology (CAT), supported by UCSF PBBR, RRP IMIA, and NIH 1S10OD028511-01 grants; Joanna Phillips of UCSF BTC Biorepository and Pathology; Jennifer Clarke of UCSF Neurological Surgery / Neuro-Oncology; Brian Shy of UCSF The Human Islet and Cellular Transplantation Facility (HICTF); David Oh and Bridget Keenan of UCSF Cancer Immunotherapy Program (CIP); and all members of the laboratory of H.O. for assistance, advice, and helpful discussions.

## Funding

This study was supported by grants from the NIH R35 1R35NS142982 (H.O.)

CIRM CLIN2-15562 (H.O.)

CRI5496 (H.O.)

Benjamin J. Cohn Fund and Esther Simon Memorial Fund through the UCSF Research

Evaluation and Allocation Committee (REAC) program (T.N.)

German Research Foundation (DFG, GA 3535/1-1) (M.G.)

ARPA-H contract D24AC00084 (W.A.L)

## Author contributions

Conceptualization: T.N., P.B.W., H.O.

Methodology: T.N., P.B.W., M.S.S., H.O.

Resources: P.B.W., M.S.S, A.Y., S.L., R.Z, W.A.L.

Investigation: T.N., P.B.W., M.S.S., A.Y., S.L., A.Z., J.L., M.G., H.L.B., R.Z., R.A.

Formal analysis: T.N., P.B.W., H.O.

Visualization: T.N., P.B.W.

Funding acquisition: T.N., H.O.

Project administration: T.N., H.O.

Supervision: H.O.

Writing (original draft): T.N., P.B.W., H.O.

Writing (review and editing): all authors.

## Competing Interests

Several patents have been filed related to this work. These include, but are not limited to, US APP # 63/464,497, 17/042,032, 17/040,476, 17/069,717, 15/831,194, 15/829,370, 15/583,658, 15/096,971, and 15/543,220. R.A. consults for the Briger Foundation for Oncology Research Inc. W.A.L. is a shareholder of Gilead Sciences, Allogene, and Intellia Therapeutics, and has consulted for Cell Design Labs, Gilead, Allogene, SciFi Foods, and Synthetic Design Lab. H.O. is on the scientific advisory boards for Neuvogen and Eureka Therapeutics. The remaining authors declare no competing interests.

### Materials and Correspondence

All data are available in the main manuscript or supplementary materials. Sequencing data and the code will be made available upon publication in a peer-reviewed journal.

## References

1. R. C. Sterner, R. M. Sterner, CAR-T cell therapy: current limitations and potential strategies. Blood Cancer J. 11, 69 (2021).

2. T. Nejo, A. Yamamichi, N. D. Almeida, Y. E. Goretsky, H. Okada, Tumor antigens in glioma. Semin. Immunol. 47, 101385 (2020).

3. J. H. Choe, P. B. Watchmaker, M. S. Simic, R. D. Gilbert, A. W. Li, N. A. Krasnow, K. M. Downey, W. Yu, D. A. Carrera, A. Celli, J. Cho, J. D. Briones, J. M. Duecker, Y. E. Goretsky, R. Dannenfelser, L. Cardarelli, O. Troyanskaya, S. S. Sidhu, K. T. Roybal, H. Okada, W. A. Lim, SynNotch-CAR T cells overcome challenges of specificity, heterogeneity, and persistence in treating glioblastoma. Sci. Transl. Med. 13 (2021), doi:10.1126/scitranslmed.abe7378.

4. M. S. Simic, P. B. Watchmaker, S. Gupta, Y. Wang, S. A. Sagan, J. Duecker, C. Shepherd, D. Diebold, P. Pineo-Cavanaugh, J. Haegelin, R. Zhu, B. Ng, W. Yu, Y. Tonai, L. Cardarelli, N. R. Reddy, S. S. Sidhu, O. Troyanskaya, S. L. Hauser, M. R. Wilson, S. S. Zamvil, H. Okada, W. A. Lim, Programming tissue-sensing T cells that deliver therapies to the brain. Science 386, eadl4237 (2024).

5. L. Morsut, K. T. Roybal, X. Xiong, R. M. Gordley, S. M. Coyle, M. Thomson, W. A. Lim, Engineering customized cell sensing and response behaviors using synthetic Notch receptors. Cell 164, 780–791 (2016).

6. Anti-EGFRvIII synNotch Receptor Induced Anti-EphA2/IL-13Ralpha2 CAR (E-SYNC) T Cells (available at https://classic.clinicaltrials.gov/ct2/show/NCT06186401).

7. S. M. Albelda, CAR T cell therapy for patients with solid tumours: key lessons to learn and unlearn. Nat. Rev. Clin. Oncol. 21, 47–66 (2024).

8. S. Park, C. E. Ho, F. Birocchi, A. N. Wolff, A. A. Bouffard, C. Kelly, D. Salas-Benito, G. Escobar, A. Mucci, T. Berger, M. V. Maus, CAR-T cell therapy targeting MUC17 in gastric tumors. J. Immunother. Cancer 13, e013120 (2025).

9. S. Huang, X. Wang, Y. Wang, Y. Wang, C. Fang, Y. Wang, S. Chen, R. Chen, T. Lei, Y. Zhang, X. Xu, Y. Li, Deciphering and advancing CAR T-cell therapy with single-cell sequencing technologies. Mol. Cancer 22, 80 (2023).

10. A. Sheih, V. Voillet, L.-A. Hanafi, H. A. DeBerg, M. Yajima, R. Hawkins, V. Gersuk, S. R. Riddell, D. G. Maloney, M. E. Wohlfahrt, D. Pande, M. R. Enstrom, H.-P. Kiem, J. E. Adair, R. Gottardo, P. S. Linsley, C. J. Turtle, Clonal kinetics and single-cell transcriptional profiling of CAR-T cells in patients undergoing CD19 CAR-T immunotherapy. Nat. Commun. 11, 219 (2020).

11. X. Li, X. Guo, Y. Zhu, G. Wei, Y. Zhang, X. Li, H. Xu, J. Cui, W. Wu, J. He, M. E. Ritchie, T. M. Weiskittel, H. Li, H. Yu, L. Ding, M. Shao, Q. Luo, X. Xu, X. Teng, A. H. Chang, J. Zhang, H. Huang, Y. Hu, Single-cell transcriptomic analysis reveals BCMA CAR-T cell dynamics in a patient with refractory primary plasma cell leukemia. Mol. Ther. 29, 645–657 (2021).

12. T. L. Wilson, H. Kim, C.-H. Chou, D. Langfitt, R. C. Mettelman, A. A. Minervina, E. K. Allen, J.-Y. Métais, M. V. Pogorelyy, J. M. Riberdy, M. P. Velasquez, P. Kottapalli, S. Trivedi, S. R. Olsen, T. Lockey, C. Willis, M. M. Meagher, B. M. Triplett, A. C. Talleur, S. Gottschalk, J. C. Crawford, P. G. Thomas, Common trajectories of highly effective CD19-specific CAR T cells identified by endogenous T-cell receptor lineages. Cancer Discov. 12, 2098–2119 (2022).

13. Z. Jackson, C. Hong, R. Schauner, B. Dropulic, P. F. Caimi, M. de Lima, M. F. Giraudo, K. Gupta, J. S. Reese, T. H. Hwang, D. N. Wald, Sequential single-cell transcriptional and protein marker profiling reveals TIGIT as a marker of CD19 CAR-T cell dysfunction in patients with non-Hodgkin lymphoma. Cancer Discov. 12, 1886–1903 (2022).

14. K. M. Dhodapkar, A. D. Cohen, A. Kaushal, A. L. Garfall, R. J. Manalo, A. R. Carr, S. S. McCachren, E. A. Stadtmauer, S. F. Lacey, J. J. Melenhorst, C. H. June, M. C. Milone, M. V. Dhodapkar, Changes in bone marrow tumor and immune cells correlate with durability of remissions following BCMA CAR T therapy in myelomaBlood Cancer Discov. 3, 490–501 (2022).

15. Z. Good, J. Y. Spiegel, B. Sahaf, M. B. Malipatlolla, Z. J. Ehlinger, S. Kurra, M. H. Desai, W. D. Reynolds, A. Wong Lin, P. Vandris, F. Wu, S. Prabhu, M. P. Hamilton, J. S. Tamaresis, P. J. Hanson, S. Patel, S. A. Feldman, M. J. Frank, J. H. Baird, L. Muffly, G. K. Claire, J. Craig, K. A. Kong, D. Wagh, J. Coller, S. C. Bendall, R. J. Tibshirani, S. K. Plevritis, D. B. Miklos, C. L. Mackall, Post-infusion CAR TReg cells identify patients resistant to CD19-CAR therapy. Nat. Med. 28, 1860–1871 (2022).

16. J. Zhang, Y. Hu, J. Yang, W. Li, M. Zhang, Q. Wang, L. Zhang, G. Wei, Y. Tian, K. Zhao, Chen, B. Tan, J. Cui, D. Li, Y. Li, Y. Qi, D. Wang, Y. Wu, D. Li, B. Du, M. Liu, H. Huang, Non-viral, specifically targeted CAR-T cells achieve high safety and efficacy in B-NHL. Nature 609, 369–374 (2022).

17. Q. Deng, G. Han, N. Puebla-Osorio, M. C. J. Ma, P. Strati, B. Chasen, E. Dai, M. Dang, N. Jain, H. Yang, Y. Wang, S. Zhang, R. Wang, R. Chen, J. Showell, S. Ghosh, S. Patchva, Q. Zhang, R. Sun, F. Hagemeister, L. Fayad, F. Samaniego, H. C. Lee, L. J. Nastoupil, N. Fowler, R. Eric Davis, J. Westin, S. S. Neelapu, L. Wang, M. R. Green, Characteristics of anti-CD19 CAR T cell infusion products associated with efficacy and toxicity in patients with large B cell lymphomas. Nat. Med. 26, 1878–1887 (2020).

18. C. A. Lareau, Y. Yin, K. Maurer, K. D. Sandor, B. Daniel, G. Yagnik, J. Peña, J. C. Crawford, A. M. Spanjaart, J. C. Gutierrez, N. J. Haradhvala, J. M. Riberdy, T. Abay, R. R. Stickels, J. M. Verboon, V. Liu, F. A. Buquicchio, F. Wang, J. Southard, R. Song, W. Li, A. Shrestha, L. Parida, G. Getz, M. V. Maus, S. Li, A. Moore, Z. J. Roberts, L. S. Ludwig, A. C. Talleur, P. G. Thomas, H. Dehghani, T. Pertel, A. Kundaje, S. Gottschalk, T. L. Roth, M. J. Kersten, C. J. Wu, R. G. Majzner, A. T. Satpathy, Latent human herpesvirus 6 is reactivated in CAR T cells. Nature 623, 608–615 (2023).

19. X. Yu, M. D. Jain, M. A. Menges, L. Cen, J. D. Noble, R. Atkins, T. J. Mohammad, C. A. Bachmeier, S. Naderinezhad, K. Reid, S. Corallo, S. J. Yoder, C. Zhang, L. Zhang, J. A. Miranda, B. Shah, J. C. Chavez, R. S. Hesterberg, L. C. Delgado, C. Savid-Frontera, P. C. Rodriguez, J. L. Cleveland, X. Wang, M. L. Davila, F. L. Locke, Comparison of axicabtagene ciloleucel and tisagenlecleucel patient CAR-T cell products by single-cell RNA sequencing. J. Immunother. Cancer 13, e011807 (2025).

20. R. G. Majzner, S. Ramakrishna, K. W. Yeom, S. Patel, H. Chinnasamy, L. M. Schultz, R. M. Richards, L. Jiang, V. Barsan, R. Mancusi, A. C. Geraghty, Z. Good, A. Y. Mochizuki, S. M. Gillespie, A. M. S. Toland, J. Mahdi, A. Reschke, E. H. Nie, I. J. Chau, M. C. Rotiroti, C. W. Mount, C. Baggott, S. Mavroukakis, E. Egeler, J. Moon, C. Erickson, S. Green, M. Kunicki, M. Fujimoto, Z. Ehlinger, W. Reynolds, S. Kurra, K. E. Warren, S. Prabhu, H. Vogel, L. Rasmussen, T. T. Cornell, S. Partap, P. G. Fisher, C. J. Campen, M. G. Filbin, G. Grant, B. Sahaf, K. L. Davis, S. A. Feldman, C. L. Mackall, M. Monje, GD2-CAR T cell therapy for H3K27M-mutated diffuse midline gliomas. Nature 603, 934–941 (2022).

21. S. J. Bagley, Z. A. Binder, L. Lamrani, E. Marinari, A. S. Desai, M. P. Nasrallah, E. Maloney, S. Brem, R. A. Lustig, G. Kurtz, M. Alonso-Basanta, P.-E. Bonté, C. Goudot, W. Richer, E. Piaggio, S. Kothari, L. Guyonnet, C. L. Guerin, J. J. Waterfall, S. Mohan, W.-T. Hwang, O. Y. Tang, M. Logun, M. Bhattacharyya, K. Markowitz, D. Delman, A. Marshall, E. J. Wherry, S. Amigorena, G. L. Beatty, J. L. Brogdon, E. Hexner, D. Migliorini, C. Alanio, D. M. O’Rourke, Repeated peripheral infusions of anti-EGFRvIII CAR T cells in combination with pembrolizumab show no efficacy in glioblastoma: a phase 1 trial. Nat Cancer 5, 517–531 (2024).

22. G. Zhong, X. Zhang, R. Zhao, Z. Guo, C. Wang, C. Yu, D. Liu, K. Hu, Y. Gao, B. Zhao, X. Liu, X. Shi, L. Chen, Y. Li, L. Yu, The high efficacy of claudin18.2-targeted CAR-T cell therapy in advanced pancreatic cancer with an antibody-dependent safety strategy. Mol. Ther. 33, 2778–2788 (2025).

23. C.-H. Li, S. Sharma, A. A. Heczey, M. L. Woods, D. H. M. Steffin, C. U. Louis, B. J. Grilley, S. G. Thakkar, M. Wu, T. Wang, C. M. Rooney, M. K. Brenner, H. E. Heslop, Long-term outcomes of GD2-directed CAR-T cell therapy in patients with neuroblastoma. Nat. Med. 31, 1125–1129 (2025).

24. M. A. Aznar, C. R. Good, J. S. Barber-Rotenberg, S. Agarwal, W. Wilson, A. Watts, Z. Zhang, D. Gonzales, G. Donahue, W.-T. Hwang, A. K. Rennels, A. J. Rech, S. Kuramitsu, H. Huang, K. M. Glastad, K. A. Alexander, G. Plesa, E. Dowd, A. Brennan, D. L. Siegel, J. Tanyi, Haas, D. A. Torigian, G. Nadolski, V. E. Gonzalez, E. O. Hexner, J. A. Fraietta, J. K. Jadlowsky, R. M. Young, S. L. Berger, C. H. June, M. H. O’Hara, Clinical and molecular dissection of CAR T cell resistance in pancreatic cancer. Cell Rep. Med. 6, 102301 (2025).

25. J. J. Melenhorst, G. M. Chen, M. Wang, D. L. Porter, C. Chen, M. A. Collins, P. Gao, S. Bandyopadhyay, H. Sun, Z. Zhao, S. Lundh, I. Pruteanu-Malinici, C. L. Nobles, S. Maji, N. V. Frey, S. I. Gill, A. W. Loren, L. Tian, I. Kulikovskaya, M. Gupta, D. E. Ambrose, M. M. Davis, J. A. Fraietta, J. L. Brogdon, R. M. Young, A. Chew, B. L. Levine, D. L. Siegel, C. Alanio, E. J. Wherry, F. D. Bushman, S. F. Lacey, K. Tan, C. H. June, Decade-long leukaemia remissions with persistence of CD4+ CAR T cells. Nature 602, 503–509 (2022).

26. N. J. Haradhvala, M. B. Leick, K. Maurer, S. H. Gohil, R. C. Larson, N. Yao, K. M. E. Gallagher, K. Katsis, M. J. Frigault, J. Southard, S. Li, M. C. Kann, H. Silva, M. Jan, K. Rhrissorrakrai, F. Utro, C. Levovitz, R. A. Jacobs, K. Slowik, B. P. Danysh, K. J. Livak, L. Parida, J. Ferry, C. Jacobson, C. J. Wu, G. Getz, M. V. Maus, Distinct cellular dynamics associated with response to CAR-T therapy for refractory B cell lymphoma. Nat. Med. 28, 1848–1859 (2022).

27. S. C. Hicks, F. W. Townes, M. Teng, R. A. Irizarry, Missing data and technical variability in single-cell RNA-sequencing experiments. Biostatistics 19, 562–578 (2018).

28. N. J. Haradhvala, M. V. Maus, Understanding mechanisms of response to CAR T-cell therapy through single-cell sequencing: Insights and challenges. Blood Cancer Discov. 5, 86– 89 (2024).

29. 10x Genomics, CG000621_RevD -Technical Note: Custom Probe Design for Visium Spatial Gene Expression and Chromium Single Cell Gene Expression Flex (10x Genomics, Pleasanton, CA, 2024; https://www.10xgenomics.com/support/flex-gene-expression/documentation/steps/experimental-design-and-planning/custom-probe-design-for-visium-spatial-gene-expression-and-chromium-single-cell-gene-expression-flex).

30. W. J. Youden, Index for rating diagnostic tests. Cancer 3, 32–35 (1950).

31. R. A. Seder, R. Ahmed, Similarities and differences in CD4+ and CD8+ effector and memory T cell generation. Nat. Immunol. 4, 835–842 (2003).

32. M. Andreatta, J. Corria-Osorio, S. Müller, R. Cubas, G. Coukos, S. J. Carmona, Interpretation of T cell states from single-cell transcriptomics data using reference atlases. Nat. Commun. 12, 2965 (2021).

33. M. Andreatta, L. Hérault, P. Gueguen, D. Gfeller, A. J. Berenstein, S. J. Carmona, Semi-supervised integration of single-cell transcriptomics data. Nat. Commun. 15, 872 (2024).

34. L. Zheng, S. Qin, W. Si, A. Wang, B. Xing, R. Gao, X. Ren, L. Wang, X. Wu, J. Zhang, N. Wu, N. Zhang, H. Zheng, H. Ouyang, K. Chen, Z. Bu, X. Hu, J. Ji, Z. Zhang, Pan-cancer single-cell landscape of tumor-infiltrating T cells. Science 374, abe6474 (2021).

35. S. Spiegel, S. Milstien, The outs and the ins of sphingosine-1-phosphate in immunity. Nat. Rev. Immunol. 11, 403–415 (2011).

36. D. Amsen, K. P. J. M. van Gisbergen, P. Hombrink, R. A. W. van Lier, Tissue-resident memory T cells at the center of immunity to solid tumors. Nat. Immunol. 19, 538–546 (2018).

37. B. V. Kumar, W. Ma, M. Miron, T. Granot, R. S. Guyer, D. J. Carpenter, T. Senda, X. Sun, S.-H. Ho, H. Lerner, A. L. Friedman, Y. Shen, D. L. Farber, Human tissue-resident memory T cells are defined by core transcriptional and functional signatures in lymphoid and mucosal sites. Cell Rep. 20, 2921–2934 (2017).

38. P. A. Szabo, H. M. Levitin, M. Miron, M. E. Snyder, T. Senda, J. Yuan, Y. L. Cheng, E. C. Bush, P. Dogra, P. Thapa, D. L. Farber, P. A. Sims, Single-cell transcriptomics of human T cells reveals tissue and activation signatures in health and disease. Nat. Commun. 10, 4706 (2019).

39. G. La Manno, R. Soldatov, A. Zeisel, E. Braun, H. Hochgerner, V. Petukhov, K. Lidschreiber, M. E. Kastriti, P. Lönnerberg, A. Furlan, J. Fan, L. E. Borm, Z. Liu, D. van Bruggen, J. Guo, X. He, R. Barker, E. Sundström, G. Castelo-Branco, P. Cramer, I. Adameyko, S. Linnarsson, P. V. Kharchenko, RNA velocity of single cells. Nature 560, 494– 498 (2018).

40. F. Odoardi, C. Sie, K. Streyl, V. K. Ulaganathan, C. Schläger, D. Lodygin, K. Heckelsmiller, W. Nietfeld, J. Ellwart, W. E. F. Klinkert, C. Lottaz, M. Nosov, V. Brinkmann, R. Spang, H. Lehrach, M. Vingron, H. Wekerle, C. Flügel-Koch, A. Flügel, T cells become licensed in the lung to enter the central nervous system. Nature 488, 675–679 (2012).

41. N. V. Gavil, K. Cheng, D. Masopust, Resident memory T cells and cancer. Immunity 57, 1734–1751 (2024).

42. P. Savas, Kathleen Cuningham Foundation Consortium for Research into Familial Breast Cancer (kConFab), B. Virassamy, C. Ye, A. Salim, C. P. Mintoff, F. Caramia, R. Salgado, D. J. Byrne, Z. L. Teo, S. Dushyanthen, A. Byrne, L. Wein, S. J. Luen, C. Poliness, S. S. Nightingale, A. S. Skandarajah, D. E. Gyorki, C. M. Thornton, P. A. Beavis, S. B. Fox, P. K. Darcy, T. P. Speed, L. K. Mackay, P. J. Neeson, S. Loi, Single-cell profiling of breast cancer T cells reveals a tissue-resident memory subset associated with improved prognosis. Nat. Med. 24, 986–993 (2018).

43. 10x Genomics, CG000478_RevD Demonstrated Protocol: Fixation of Cells & Nuclei for Chromium Fixed RNA Profiling (10x Genomics, Pleasanton, CA, 2024; https://www.10xgenomics.com/support/flex-gene-expression/documentation/steps/sample-prep/fixation-of-cells-and-nuclei-for-chromium-single-cell-gene-expression-flex).

44. 10x Genomics, *CG000527_RevF User Guide: Chromium fixed RNA profiling reagent kits for multiplexed samples* (10x Genomics, Pleasanton, CA, 2024).

45. Y. Hao, S. Hao, E. Andersen-Nissen, W. M. Mauck 3rd, S. Zheng, A. Butler, M. J. Lee, A. J. Wilk, C. Darby, M. Zager, P. Hoffman, M. Stoeckius, E. Papalexi, E. P. Mimitou, J. Jain, A. Srivastava, T. Stuart, L. M. Fleming, B. Yeung, A. J. Rogers, J. M. McElrath, C. A. Blish, R. Gottardo, P. Smibert, R. Satija, Integrated analysis of multimodal single-cell data. Cell 184, 3573–3587.e29 (2021).

46. P.-L. Germain, A. Lun, C. Garcia Meixide, W. Macnair, M. D. Robinson, Doublet identification in single-cell sequencing data using scDblFinder. F1000Res. 10, 979 (2021).

47. T. Stuart, A. Butler, P. Hoffman, C. Hafemeister, E. Papalexi, W. M. Mauck 3rd, Y. Hao, M. Stoeckius, P. Smibert, R. Satija, Comprehensive Integration of Single-Cell Data. Cell 177, 1888–1902.e21 (2019).

48. M. Andreatta, S. J. Carmona, UCell: Robust and scalable single-cell gene signature scoring. Comput. Struct. Biotechnol. J. 19, 3796–3798 (2021).

49. Y. Chu, E. Dai, Y. Li, G. Han, G. Pei, D. R. Ingram, K. Thakkar, J.-J. Qin, M. Dang, X. Le, C. Hu, Q. Deng, A. Sinjab, P. Gupta, R. Wang, D. Hao, F. Peng, X. Yan, Y. Liu, S. Song, S. Zhang, J. V. Heymach, A. Reuben, Y. Y. Elamin, M. P. Pizzi, Y. Lu, R. Lazcano, J. Hu, M. Li, M. Curran, A. Futreal, A. Maitra, A. A. Jazaeri, J. A. Ajani, C. Swanton, X.-D. Cheng, H. A. Abbas, M. Gillison, K. Bhat, A. J. Lazar, M. Green, K. Litchfield, H. Kadara, C. Yee, L. Wang, Pan-cancer T cell atlas links a cellular stress response state to immunotherapy resistance. Nat. Med. 29, 1550–1562 (2023).

