## Supplementary Figures for "Spike-in probe-enhanced single-cell RNA-seq reveals post-infusion transcriptomic remodeling of “prime-and-kill” synNotch-CAR-T cells"

Takahide Nejo et al.

**The PDF file includes:**

Figs. S1 to S17

**Other supplementary materials for this manuscript include:**

Supplementary Tables

S1. Spike-in probes (Related to Fig. 1).

S2. Single-cell RNA-seq quality control values (Related to Fig. 2).

S3. Differential gene expression (DGE) analysis of $CD8^+$ synNotch-positive cells comparing six cell clusters (Related to Fig. 3G).

S4. DGE analysis of $CD4^+$ synNotch-positive cells comparing four cell clusters (Related to Fig. S7C).

S5. DGE analysis of $CD8^+$ synNotch-positive cells comparing three *in vivo* sample types (Related to Fig. 4D).

S6. DGE analysis of $CD8^+$ synNotch-positive cells comparing pre-infusion and brain samples (Related to Fig. 4D).

S7. DGE analysis of $CD4^+$ synNotch-positive cells comparing three *in vivo* sample types (Related to Fig. S9D).

S8. DGE analysis of $CD4^+$ synNotch-positive cells comparing pre-infusion and brain samples (Related to Fig. S9D).

S9. Pseudobulk DGE analysis of $CD8^+$ brain-derived cells comparing synNotch-positive and -negative subsets (Related to Fig. 5C–D).

S10. Pseudobulk DGE analysis of $CD4^+$ brain-derived cells comparing synNotch-positive and -negative subsets (Related to Fig. S12C–D).
S11. Pseudobulk DGE analysis of CD8+ brain-derived synNotch-positive cells comparing CAR-high and CAR-low subsets (Related to Fig. 6).
S12. Pseudobulk DGE analysis of CD4+ brain-derived synNotch-positive cells comparing CAR-high and CAR-low subsets (Related to Fig. S15).
S13. Bulk RNAseq gene expression data of the time-course co-culture experiment of B-SYNC cells with GBM6 (Related to Fig. 6L).

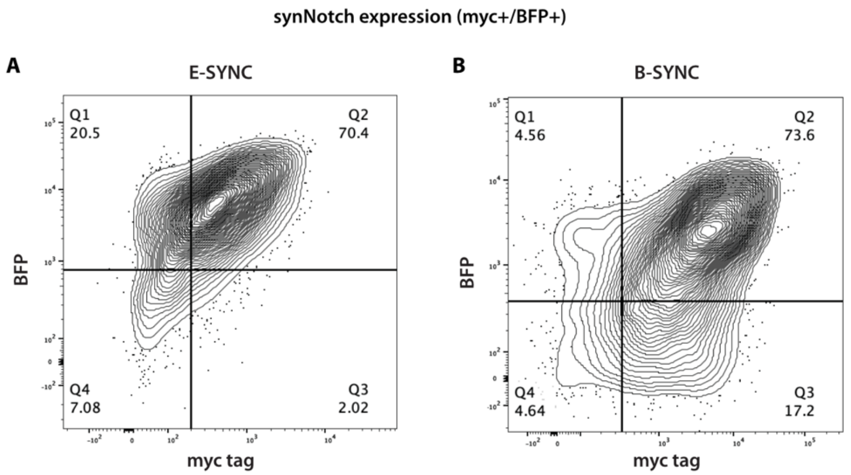


**Fig. S1. Flow cytometry analysis of synNotch expression.**

(**A–B**) SynNotch gene silencing data for E-SYNC (**A**) and B-SYNC (**B**) cells from batches independent of those used for the scRNA-seq experiment. The $BFP^+/Myc^+$ double-positive population defines synNotch-expressing cells.

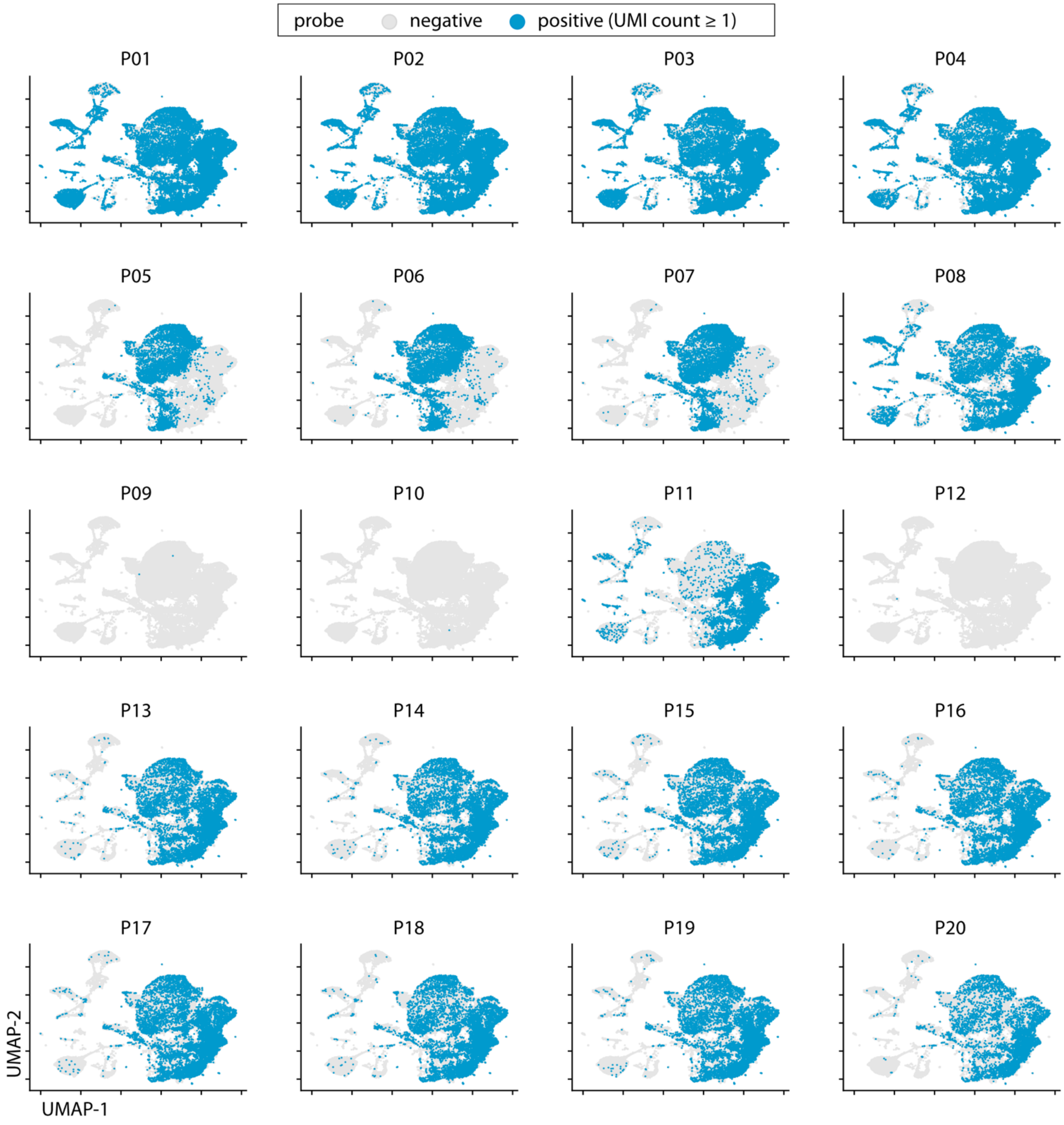


**Fig. S2. Spike-in probe positivity across the full dataset (combined mouse–human dataset).**

UMAP plots highlighting cells with unique molecular identifier (UMI) counts ≥ 1 (shown in blue) for each spike-in probe across the full dataset (n = 71,541 cells).

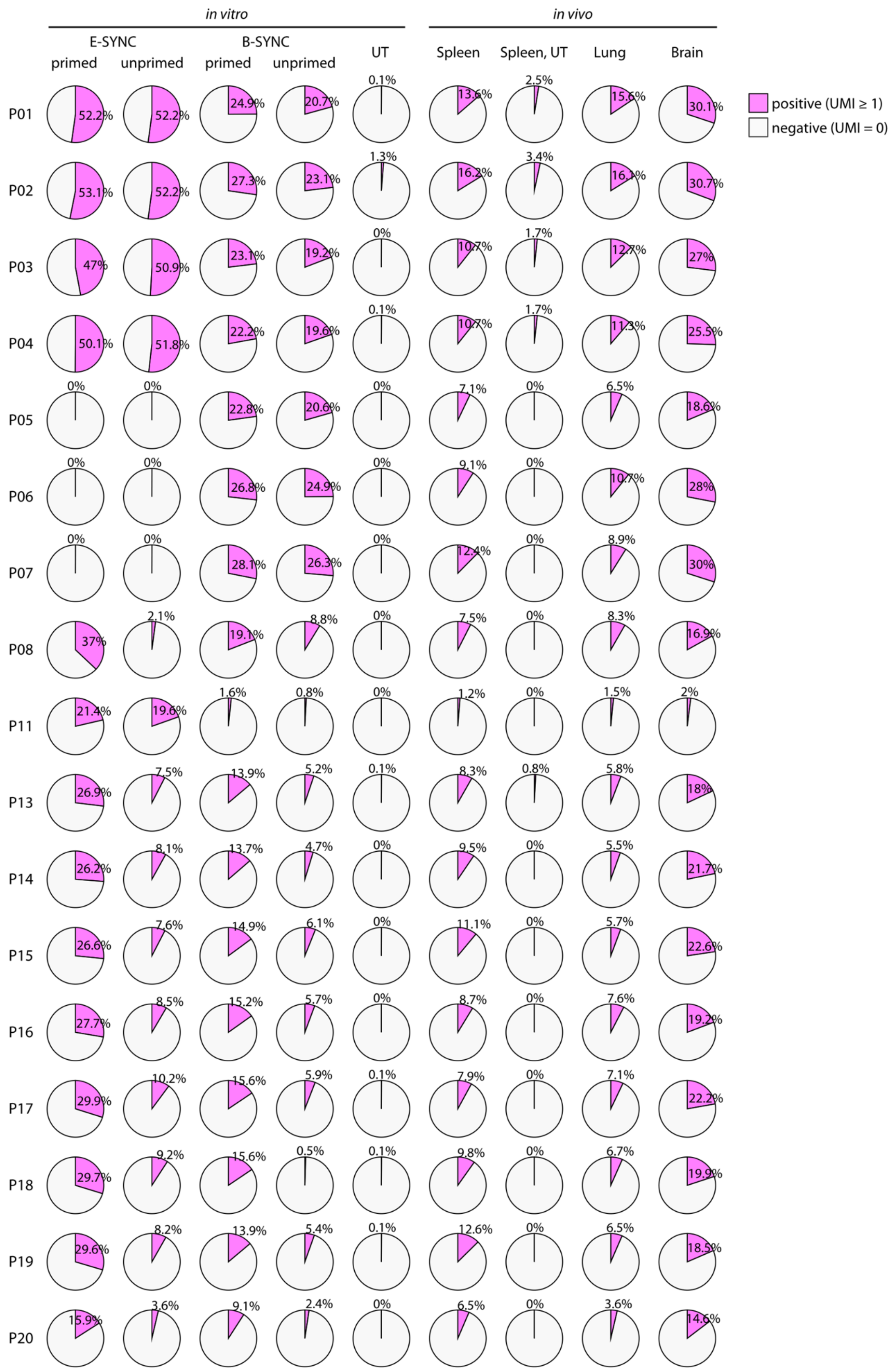
in vitro
in vivo
E-SYNC
B-SYNC
primed
unprimed
primed
unprimed
UT
Spleen
Spleen, UT
Lung
Brain
positive (UMI ≥ 1)
negative (UMI = 0)
P01 52.2% 52.2% 24.9% 20.7% 0.1% 13.6% 2.5% 15.6% 30.1%
P02 53.1% 52.2% 27.3% 23.1% 1.3% 16.2% 3.4% 16.1% 30.7%
P03 47% 50.9% 23.1% 19.2% 0% 10.7% 1.7% 12.7% 27%
P04 50.1% 51.8% 22.2% 19.6% 0.1% 10.7% 1.7% 11.3% 25.5%
P05 0% 0% 22.8% 20.6% 0% 7.1% 0% 6.5% 18.6%
P06 0% 0% 26.8% 24.9% 0% 9.1% 0% 10.7% 28%
P07 0% 0% 28.1% 26.3% 0% 12.4% 0% 8.9% 30%
P08 37% 2.1% 19.1% 8.8% 0% 7.5% 0% 8.3% 16.9%
P11 21.4% 19.6% 1.6% 0.8% 0% 1.2% 0% 1.5% 2%
P13 26.9% 7.5% 13.9% 5.2% 0.1% 8.3% 0.8% 5.8% 18%
P14 26.2% 8.1% 13.7% 4.7% 0% 9.5% 0% 5.5% 21.7%
P15 26.6% 7.6% 14.9% 6.1% 0% 11.1% 0% 5.7% 22.6%
P16 27.7% 8.5% 15.2% 5.7% 0% 8.7% 0% 7.6% 19.2%
P17 29.9% 10.2% 15.6% 5.9% 0.1% 7.9% 0% 7.1% 22.2%
P18 29.7% 9.2% 15.6% 0.5% 0.1% 9.8% 0% 6.7% 19.9%
P19 29.6% 8.2% 13.9% 5.4% 0.1% 12.6% 0% 6.5% 18.5%
P20 15.9% 3.6% 9.1% 2.4% 0% 6.5% 0% 3.6% 14.6%

**Fig. S3. Detection performance of 17 functional spike-in probes.**

Pie charts showing the proportion of human cells positive for each probe. Positivity was defined as UMI count ≥ 1.

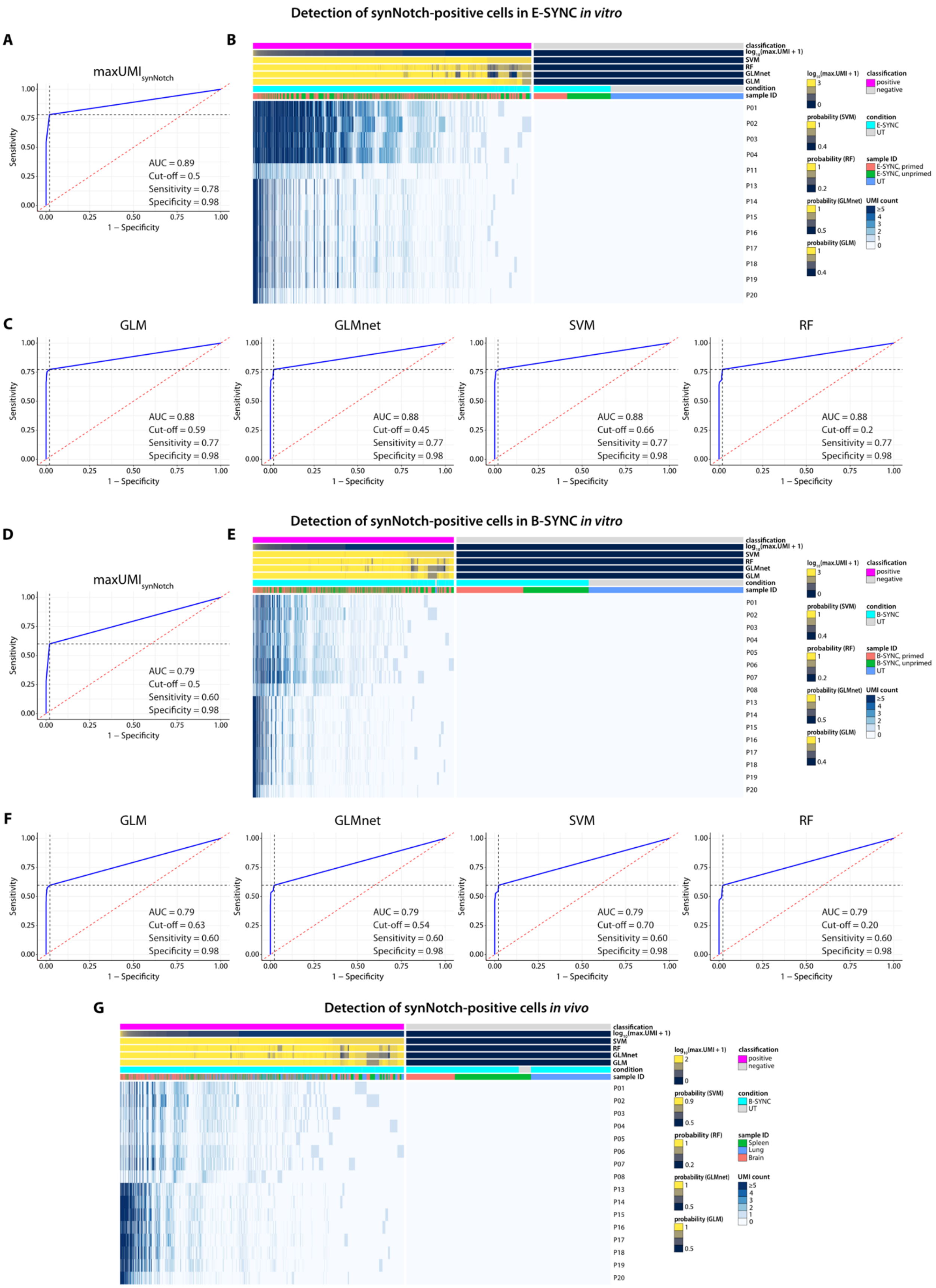

Detection of synNotch-positive cells in E-SYNC in vitro
A
maxUMI synNotch
AUC = 0.89
Cut-off = 0.5
Sensitivity = 0.78
Specificity = 0.98
Sensitivity
1 – Specificity
B
classification
log10(max.UMI + 1)
SVM
RF
GLMnet
GLM
condition
sample ID
P01
P02
P03
P04
P11
P13
P14
P15
P16
P17
P18
P19
P20
positive
negative
E-SYNC
UT
E-SYNC, primed
E-SYNC, unprimed
probability (SVM)
probability (RF)
probability (GLMnet)
probability (GLM)
UMI count
C
GLM
AUC = 0.88
Cut-off = 0.59
Sensitivity = 0.77
Specificity = 0.98
GLMnet
AUC = 0.88
Cut-off = 0.45
Sensitivity = 0.77
Specificity = 0.98
SVM
AUC = 0.88
Cut-off = 0.66
Sensitivity = 0.77
Specificity = 0.98
RF
AUC = 0.88
Cut-off = 0.2
Sensitivity = 0.77
Specificity = 0.98
Detection of synNotch-positive cells in B-SYNC in vitro
D
maxUMI synNotch
AUC = 0.79
Cut-off = 0.5
Sensitivity = 0.60
Specificity = 0.98
E
P01
P02
P03
P04
P05
P06
P07
P08
P13
P14
P15
P16
P17
P18
P19
P20
B-SYNC
UT
B-SYNC, primed
B-SYNC, unprimed
F
GLM
AUC = 0.79
Cut-off = 0.63
Sensitivity = 0.60
Specificity = 0.98
GLMnet
AUC = 0.79
Cut-off = 0.54
Sensitivity = 0.60
Specificity = 0.98
SVM
AUC = 0.79
Cut-off = 0.70
Sensitivity = 0.60
Specificity = 0.98
RF
AUC = 0.79
Cut-off = 0.20
Sensitivity = 0.60
Specificity = 0.98
G
Detection of synNotch-positive cells in vivo
B-SYNC
UT
Spleen
Lung
Brain

**Fig. S4. Machine learning-assisted classification of synNotch-positive and -negative cells.**

(**A–C**) Receiver operating characteristic (ROC) curves (**A, C**) and heatmap (**B**) showing classifier performance for detecting synNotch-positive cells in E-SYNC and untransduced (UT) cells.

(**D–F**) ROC curves (**D, F**) and heatmap (**E**) showing classifier performance for detecting synNotch-positive cells in B-SYNC and UT cells.

(**G**) Heatmap showing classifier performance for detecting synNotch-positive cells in *in vivo* samples using B-SYNC-derived classifiers.

AUC, area under the curve.

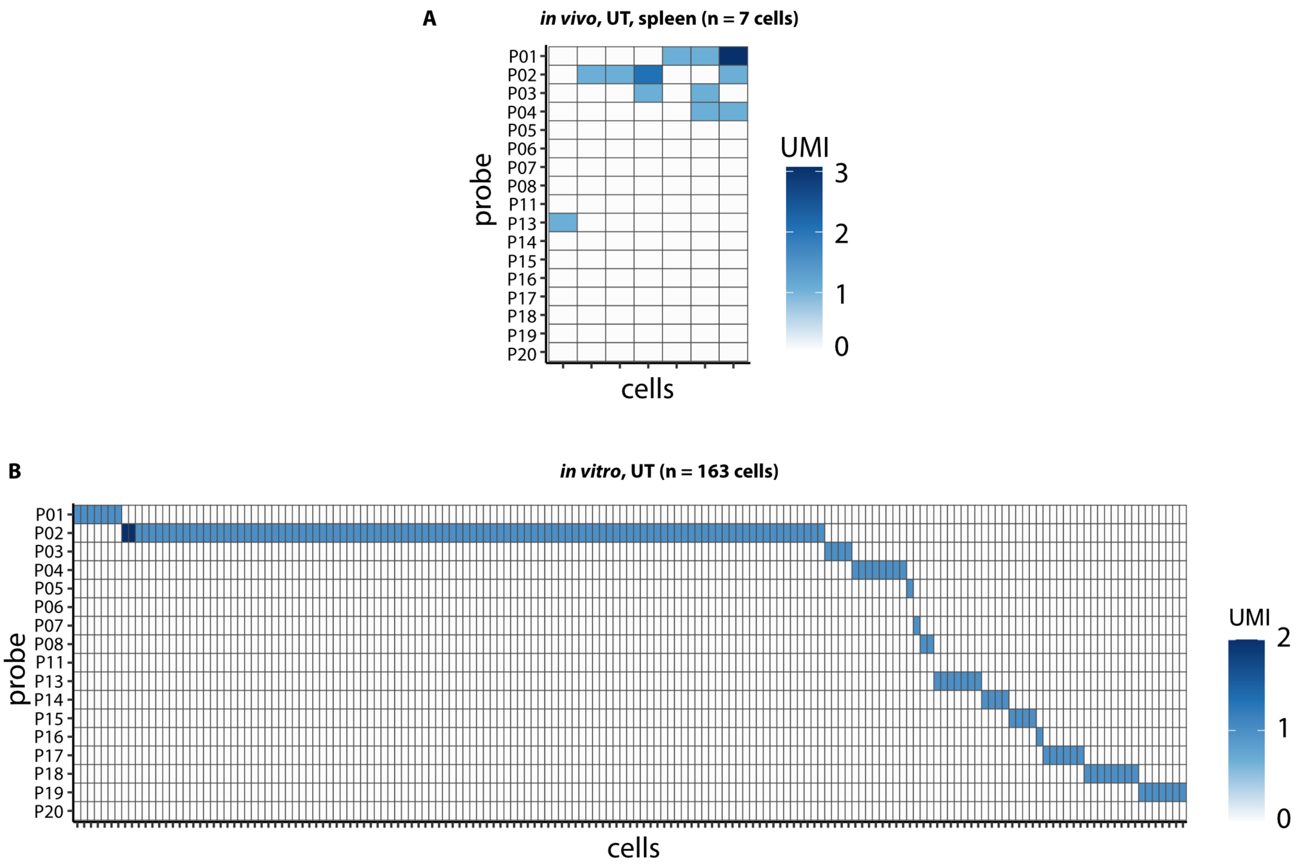


**Fig. S5. Assessment of false-positive synNotch-CAR transcript detection.**

(**A–B**) Heatmaps showing probe-level signal distributions in cells misclassified as synNotch-positive in negative controls: 7 of 119 cells (5.9%) from the spleen of a UT-treated mouse (**A**) and 163 of 8,170 cells (2.0%) from *in vitro* UT samples (**B**).

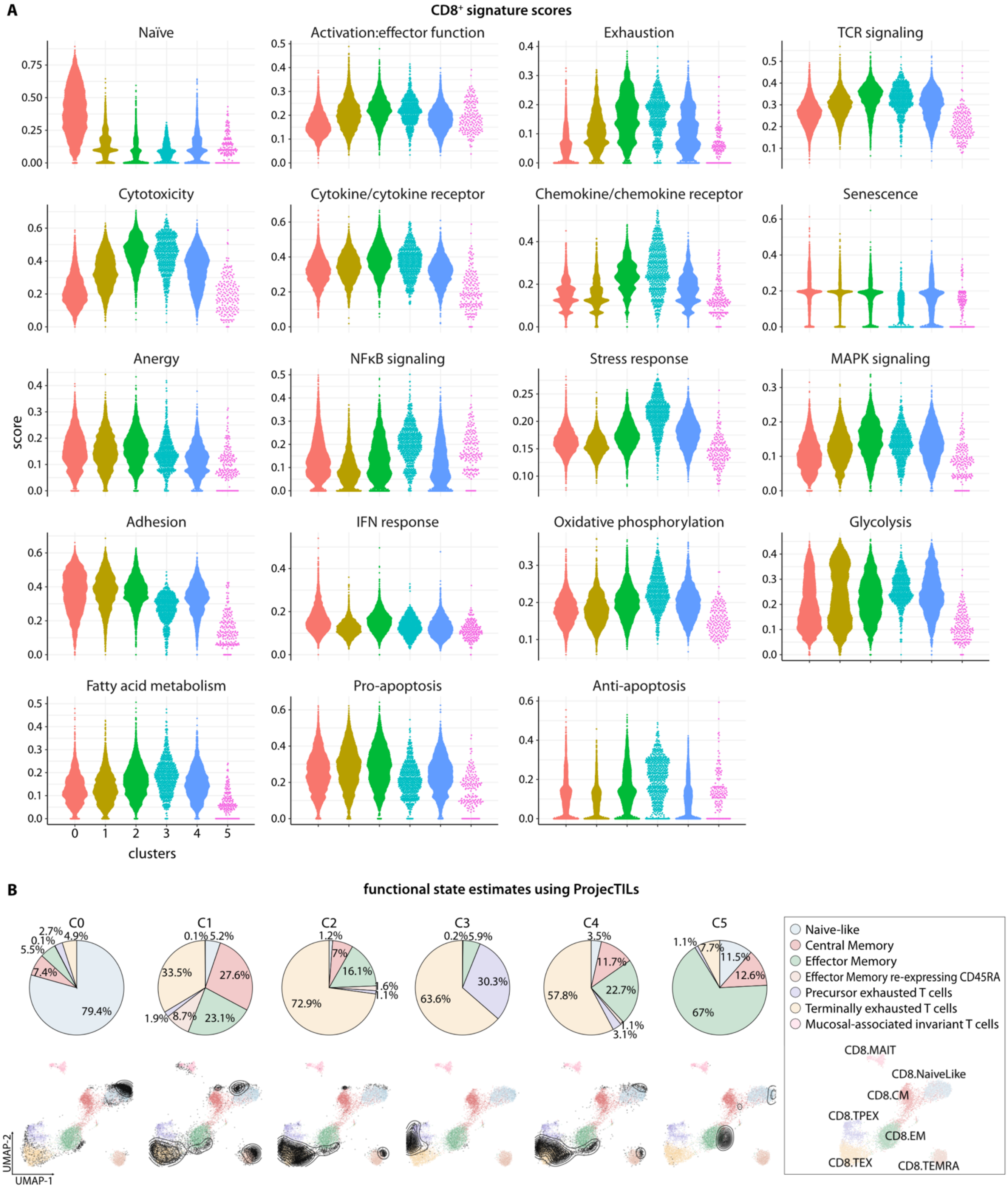
A
CD8+ signature scores
Naïve
Activation:effector function
Exhaustion
TCR signaling
Cytotoxicity
Cytokine/cytokine receptor
Chemokine/chemokine receptor
Senescence
Anergy
NFκB signaling
Stress response
MAPK signaling
Adhesion
IFN response
Oxidative phosphorylation
Glycolysis
Fatty acid metabolism
Pro-apoptosis
Anti-apoptosis
score
0 1 2 3 4 5
clusters
B
functional state estimates using ProjecTILs
C0
C1
C2
C3
C4
C5
Naive-like
Central Memory
Effector Memory
Effector Memory re-expressing CD45RA
Precursor exhausted T cells
Terminally exhausted T cells
Mucosal-associated invariant T cells
CD8.MAIT
CD8.NaiveLike
CD8.CM
CD8.TPEX
CD8.EM
CD8.TEX
CD8.TEMRA
UMAP-2
UMAP-1

**Fig. S6. CD8$^+$ T cell gene signature scores and ProjecTILs analysis results (Related to Fig. 3H).**

(**A**) Swarm plots showing the distribution of 19 CD8$^+$ signature scores across clusters *(1)*.

(**B**) Pie charts and UMAP contour plots depicting the projection of CD8$^+$ T cell clusters onto a reference human CD8$^+$ tumor-infiltrating lymphocyte (TIL) dataset *(2, 3)*.

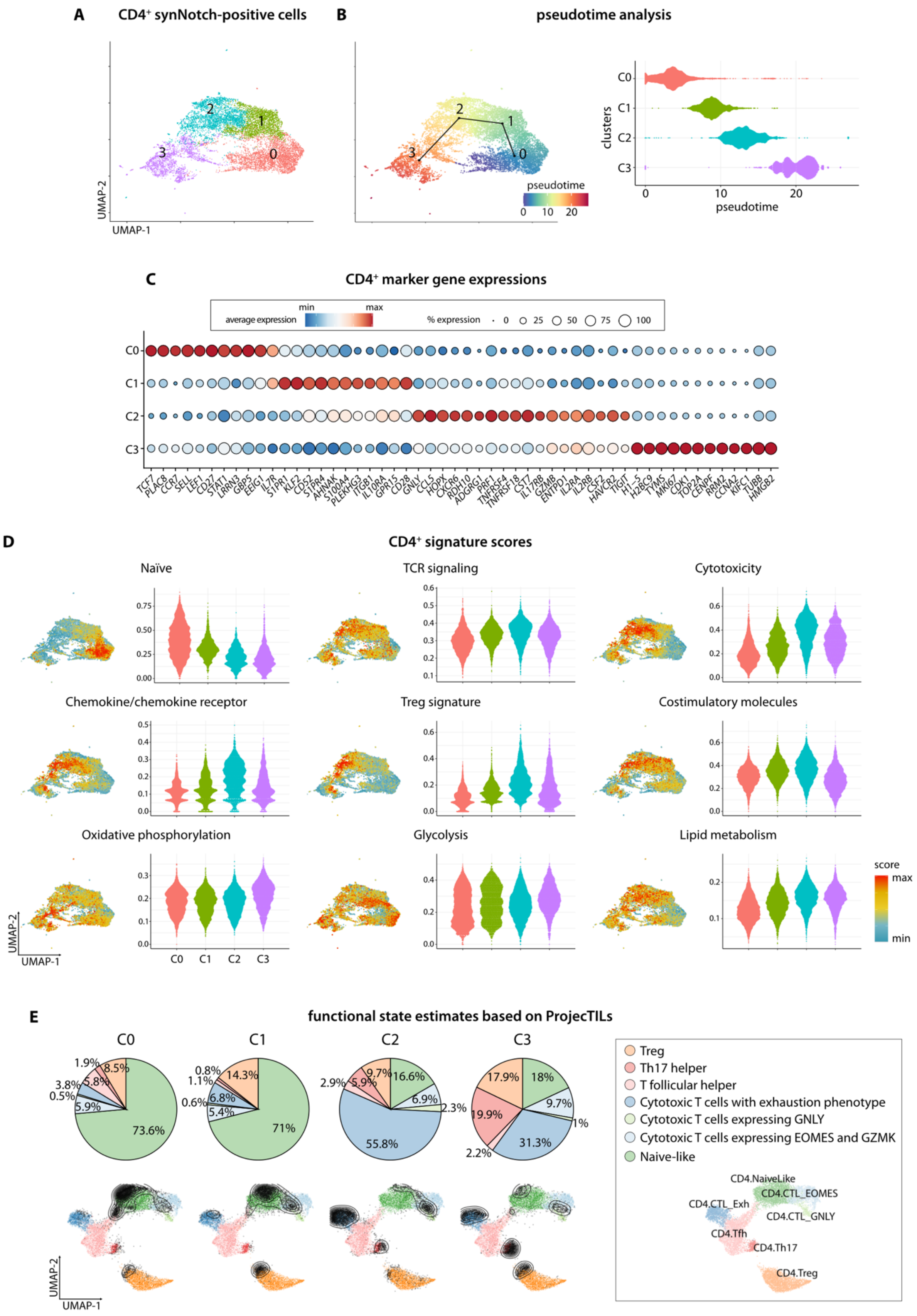
A
CD4+ synNotch-positive cells
UMAP-1
UMAP-2
B
pseudotime analysis
pseudotime
clusters
C0
C1
C2
C3
C
CD4+ marker gene expressions
average expression
min
max
% expression
0
25
50
75
100
TCF7 PLAC8 CCR7 SELL LEF1 CD27 STAT1 LRRN3 GBP5 EEIG1 IL7R S1PR1 KLF2 CD52 S1PR4 AHNAK S100A4 PLEKHG3 ITGB1 IL10RA GPR15 CD28 GNLY CCL5 HOPX CXCR6 RDH10 ADGRG1 PRF1 TNFRSF4 TNFRSF18 CST7 IL17RB GZMB ENTPD1 IL2RA IL2RB CSF2 HAVCR2 TIGIT H1-5 H2BC9 TYMS MKI67 CDK1 TOP2A CENPF RRM2 CCNA2 KIFC1 TUBB HMGB2
D
CD4+ signature scores
Naïve
TCR signaling
Cytotoxicity
Chemokine/chemokine receptor
Treg signature
Costimulatory molecules
Oxidative phosphorylation
Glycolysis
Lipid metabolism
score
max
min
E
functional state estimates based on ProjecTILs
Treg
Th17 helper
T follicular helper
Cytotoxic T cells with exhaustion phenotype
Cytotoxic T cells expressing GNLY
Cytotoxic T cells expressing EOMES and GZMK
Naive-like
CD4.NaiveLike
CD4.CTL_EOMES
CD4.CTL_Exh
CD4.CTL_GNLY
CD4.Tfh
CD4.Th17
CD4.Treg

**Fig. S7. Profiling the synNotch-positive $CD4^+$ population.**

(**A**) UMAP plot of synNotch-positive $CD4^+$ cells colored by cluster identity.

(**B**) UMAP feature plot and corresponding swarm plot displaying pseudotime estimates (slingshot) for individual cells and clusters.

(**C**) Dot plot showing curated differentially expressed genes (DEGs) upregulated in each cluster. The complete DEG list is provided in **Supplementary Table S4**.

(**D**) Swarm plots and UMAP feature plots illustrating representative $CD4^+$ signature score distributions.

(**E**) Pie charts and UMAP contour plots depicting the projection of $CD4^+$ T cell clusters onto a reference human $CD4^+$ TIL dataset *(2, 4)*.

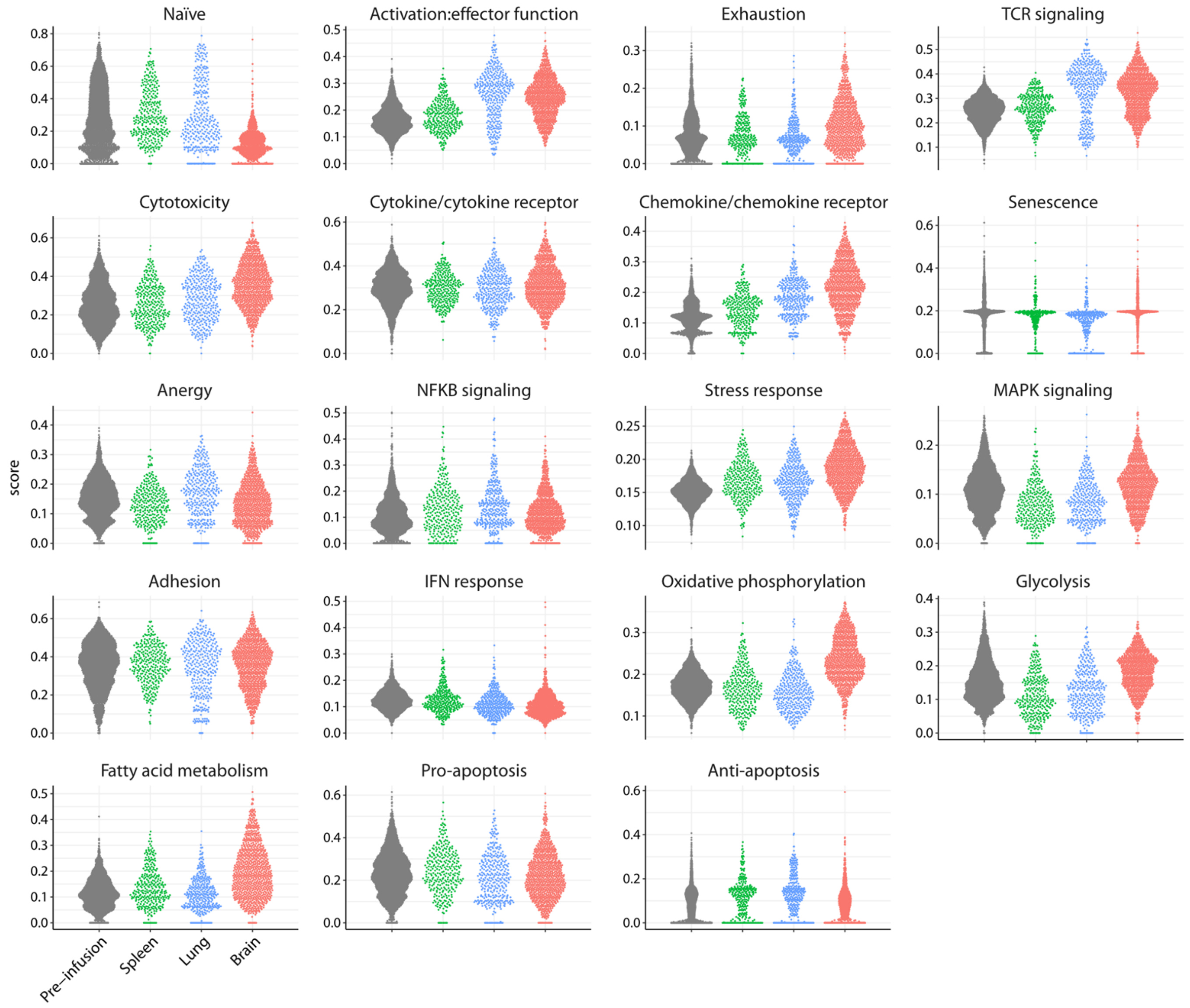


**Fig. S8. CD8⁺ T cell gene signature scores across sample types (Related to Fig. 4C).** Swarm plots showing the distribution of 19 CD8⁺ T cell gene signature scores across sample types (pre-infusion, spleen, lung, and brain) *(4)*.

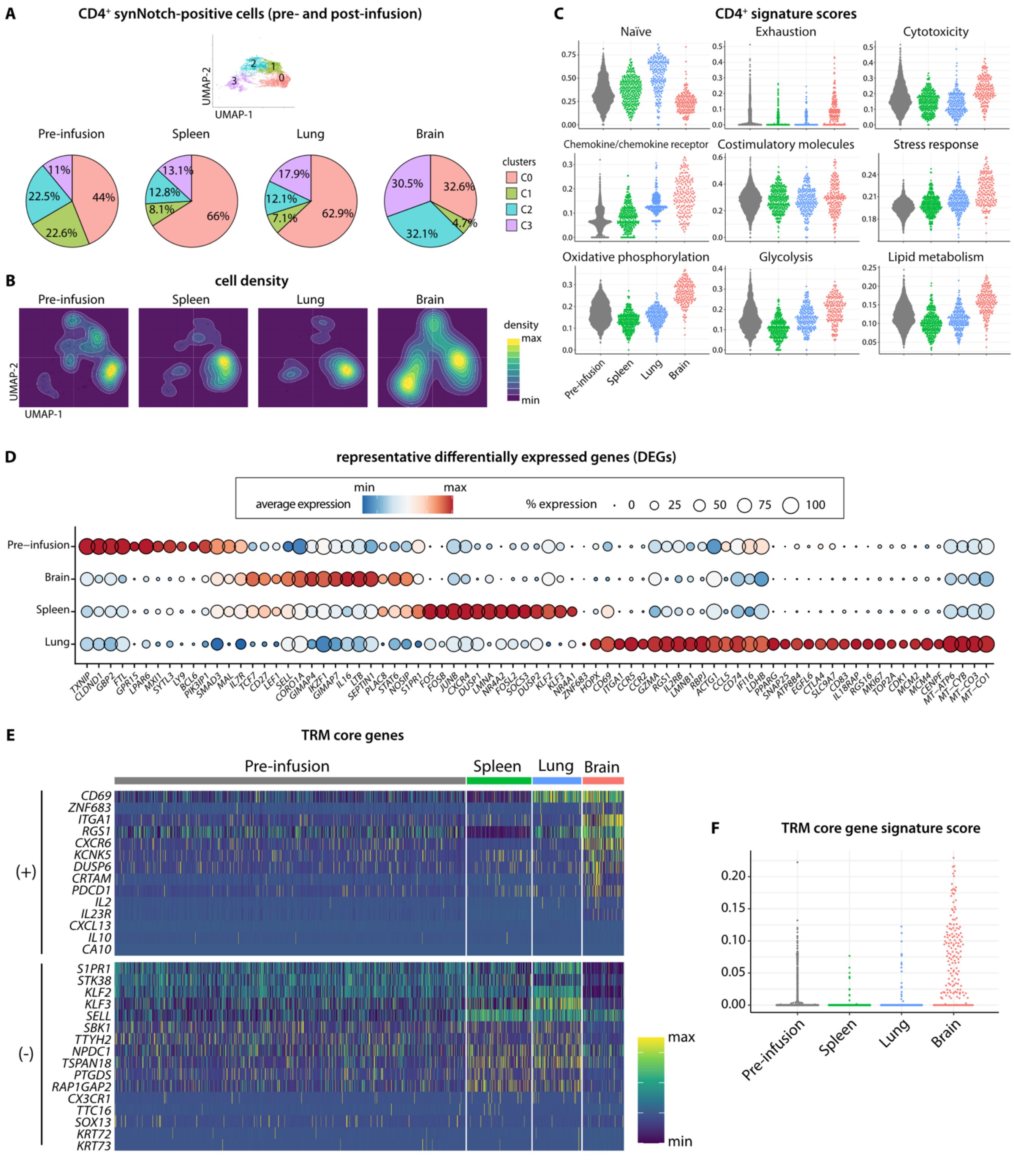


**Fig. S9.** ***In vivo*** **transcriptomic dynamics of synNotch-positive CD4+ T cells (Related to Fig. 4).**

(**A**) Pie charts comparing $CD4^+$ cluster proportions across pre-infusion (B-SYNC, unprimed), spleen, lung, and brain samples.

(**B**) UMAP density contour plots showing cell distributions for each sample type, with higher-density areas indicated by brighter colors.

(**C**) Swarm plots showing representative $CD4^+$ T cell signature score distributions across sample types.

(**D**) Dot plot showing curated differentially expressed genes (DEGs) across sample types. The complete DEG list is provided in **Supplementary Tables S7–S8**.

(**E**) Heatmap showing expression of the tissue-resident memory (TRM) core genes *(5)* in individual cells.

(**F**) Swarm plot depicting the distribution of TRM core gene signature scores. Differences among sample types were evaluated using the Kruskal–Wallis test ($P = 7.8\times10^{-117}$).

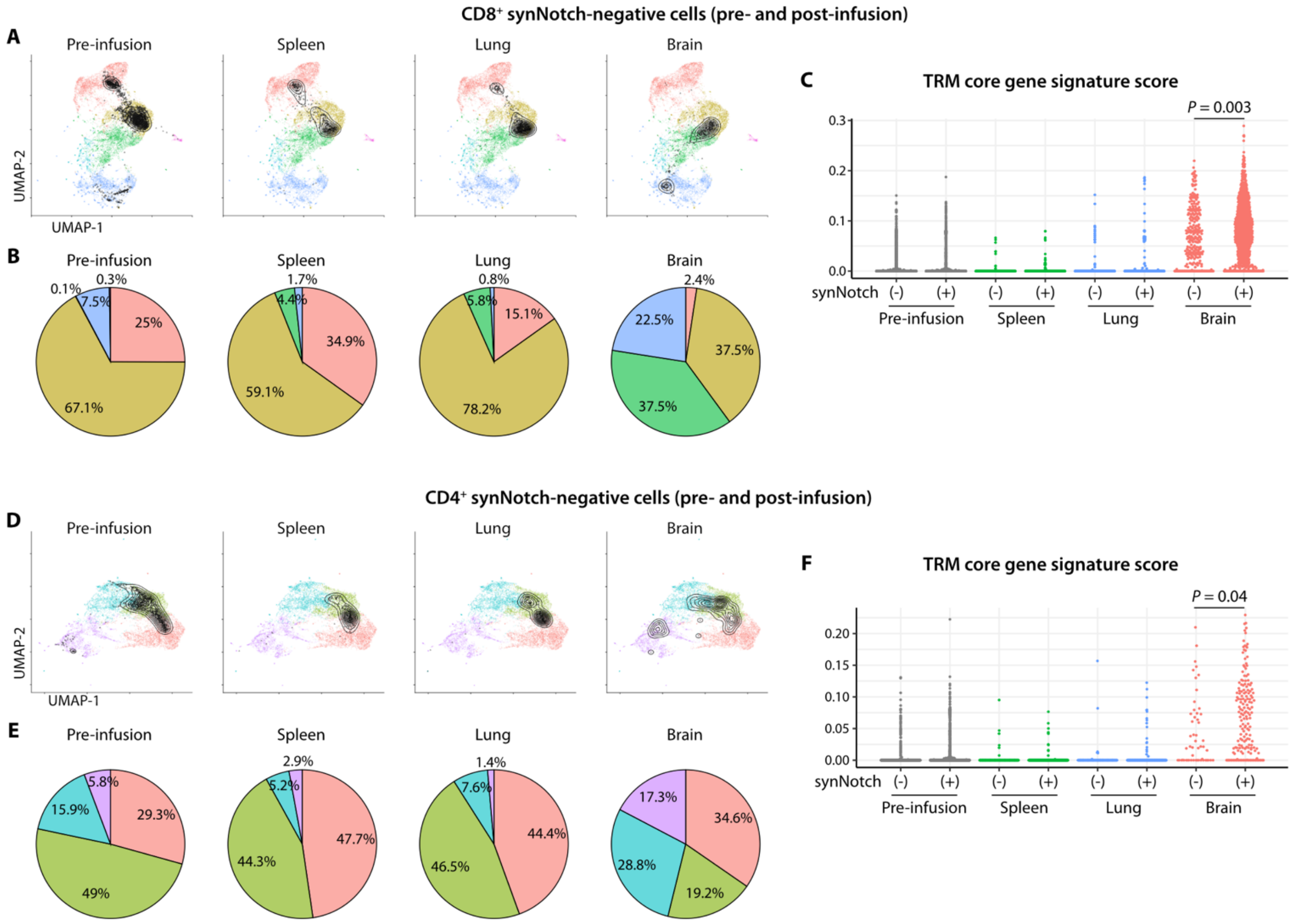


**Fig. S10. Comparison of synNotch-positive and -negative CD8⁺ and CD4⁺ cells.**

(**A–B**) UMAP contour plots (**A**) and pie charts (**B**) showing projection of synNotch-negative CD8⁺ T cells from each sample type onto the synNotch-positive CD8⁺ reference map.

(**C**) Swarm plot depicting the distribution of TRM core gene signature scores *(5)* in CD8⁺ T cells. The difference between brain-derived synNotch-positive and -negative cells was evaluated using the Kolmogorov–Smirnov test ($P = 0.003$, $D = 0.12$).

(**D–E**) UMAP contour plots (**D**) and pie charts (**E**) showing projection of synNotch-negative CD4⁺ T cells from each sample type onto the synNotch-positive CD4⁺ reference map.

(**F**) Swarm plot depicting the distribution of TRM core gene signature scores in CD4⁺ T cells. The difference between brain-derived synNotch-positive and -negative cells was evaluated using the Kolmogorov–Smirnov test ($P = 0.04$, $D = 0.22$).

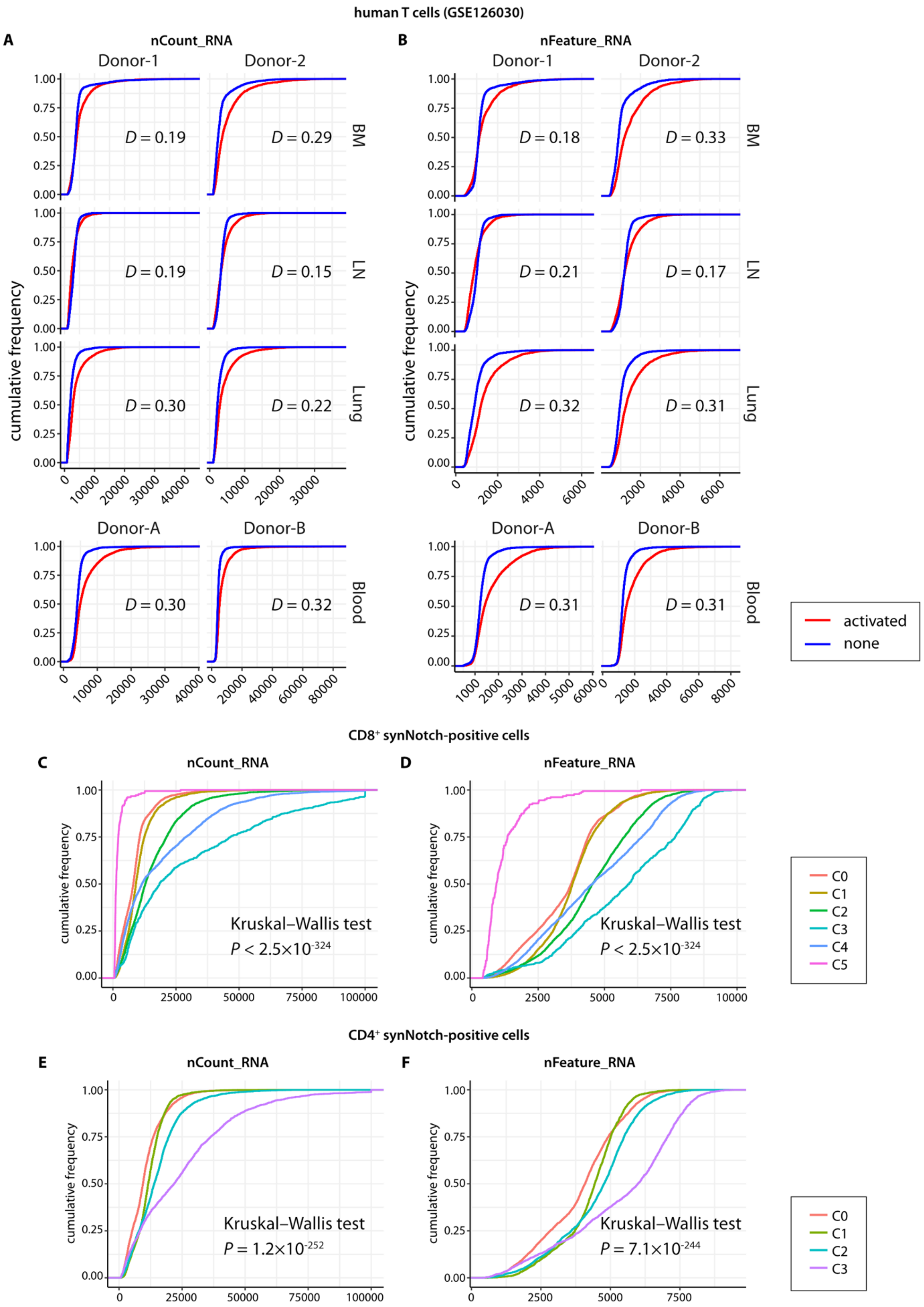

human T cells (GSE126030)
A
nCount_RNA
B
nFeature_RNA
Donor-1
Donor-2
Donor-A
Donor-B
BM
LN
Lung
Blood
cumulative frequency
D = 0.19
D = 0.29
D = 0.19
D = 0.15
D = 0.30
D = 0.22
D = 0.30
D = 0.32
D = 0.18
D = 0.33
D = 0.21
D = 0.17
D = 0.32
D = 0.31
D = 0.31
D = 0.31
activated
none
CD8+ synNotch-positive cells
C
nCount_RNA
D
nFeature_RNA
Kruskal–Wallis test
P < 2.5×10-324
Kruskal–Wallis test
P < 2.5×10-324
C0
C1
C2
C3
C4
C5
CD4+ synNotch-positive cells
E
nCount_RNA
F
nFeature_RNA
Kruskal–Wallis test
P = 1.2×10-252
Kruskal–Wallis test
P = 7.1×10-244
C0
C1
C2
C3

**Fig. S11. T cell activation is associated with increased transcriptional activity.**

(**A–F**) Empirical cumulative distribution function (ECDF) plots of nCount_RNA (total UMI counts per cell) (**A, C, E**) and nFeature_RNA (number of detected genes per cell) (**B, D, F**). Panels (A–B) show comparisons of activated and resting human T cells from multiple sources in a public dataset (GSE126030) *(5)*. Panes (C–F) show the corresponding distributions in $CD8^+$ (**C, D**) and $CD4^+$ (**E, F**) synNotch-positive populations across clusters. In panels (**C**) and (**E**), values exceeding 100,000 were capped at 100,000 for visualization. Statistical significance was determined using the Kolmogorov–Smirnov test (**A–B**) and the Kruskal–Wallis test (**C–F**). For all comparisons in panels (**A–B**), $P$ values were $< 2.2\times10^{-324}$, and D values are shown in each plot.

BM, bone marrow; LN, lymph node.

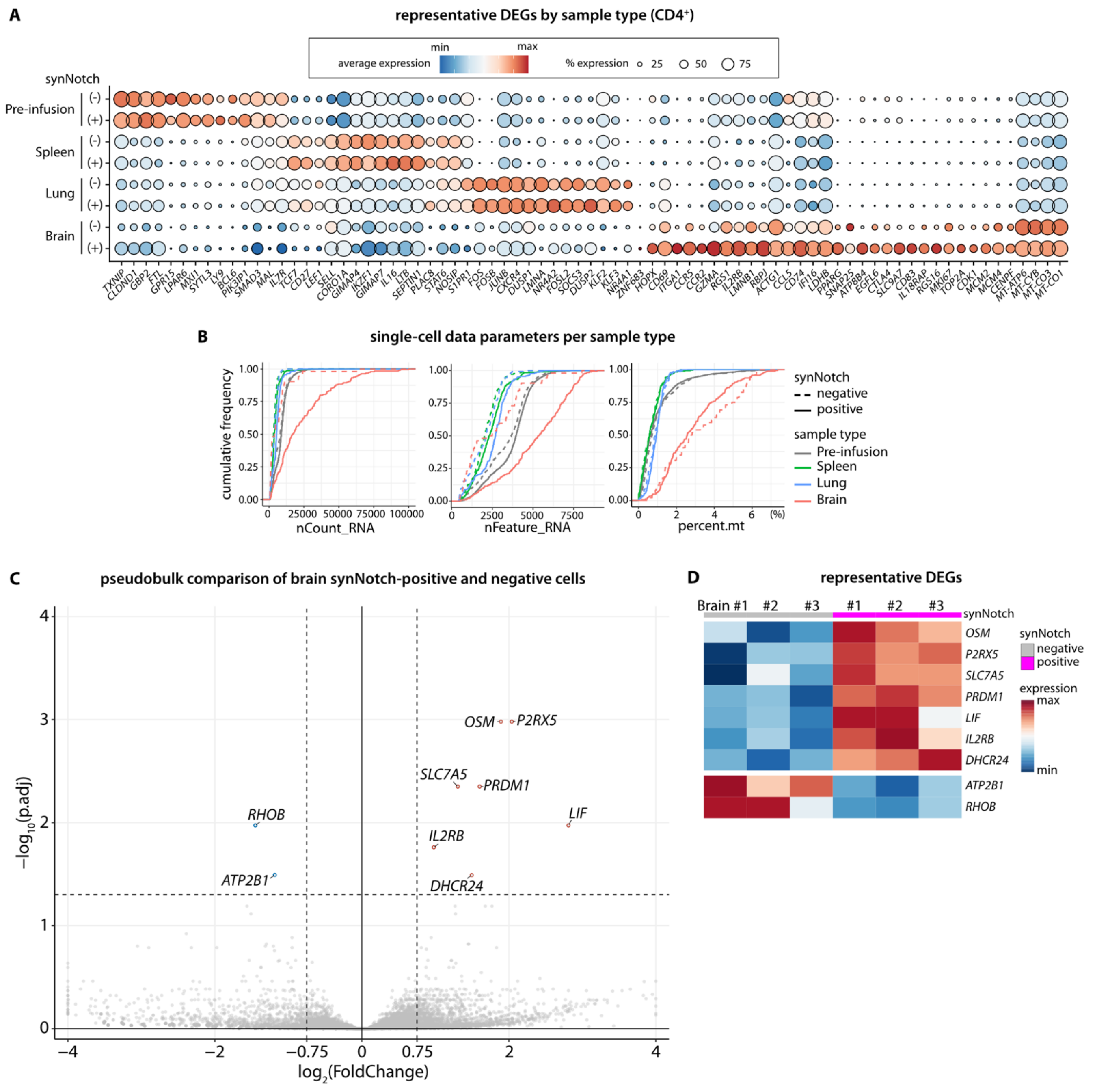


**Fig. S12. Transcriptomic remodeling by environmental signals and synNotch-mediated antigen recognition in the CD4⁺ population (Related to Fig. 5).**

(**A**) Dot plot showing the expression of the same gene list as in **Fig. S9D** across sample types, stratified by synNotch-positive and -negative cells.

(**B**) Empirical cumulative distribution function (ECDF) plots showing distributions of nCount_RNA (transcript abundance), nFeature_RNA (number of expressed genes), and percent.mt (mitochondrial transcript fraction) across sample types.

(**C, D**) Volcano plot and heatmap summarizing pseudobulk differential expression analysis comparing brain-derived $CD4^+$ synNotch-positive and -negative cells. Genes with an adjusted *P* value < 0.05 and $|\log_2(\text{Fold Change})| > 0.75$ were defined as differentially expressed genes (DEGs). DEGs are highlighted in red or blue and labeled in the volcano plot (**C**). Scaled expression levels of these DEGs are shown in the heatmap (**D**). The complete DEG list is provided in **Supplementary Table S10**.

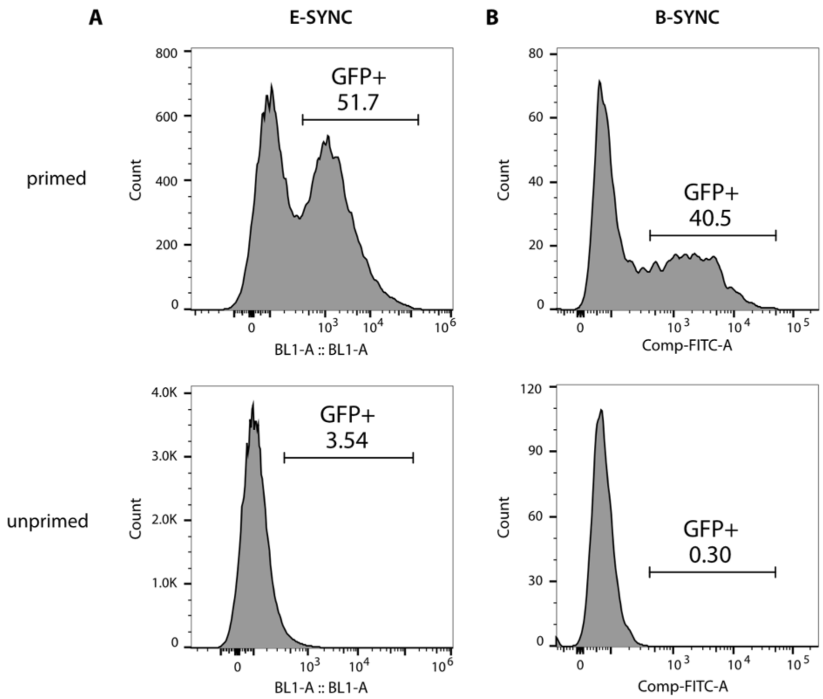


**Fig. S13. Flow cytometry analysis of CAR expression.**

(**A–B**) Flow cytometry histograms showing the proportion of CAR-expressing cells, as indicated by GFP positivity, within the live, singlet, BFP$^{+}$ positive population of *in vitro* E-SYNC (**A**) and B-SYNC (**B**) primed and unprimed conditions.

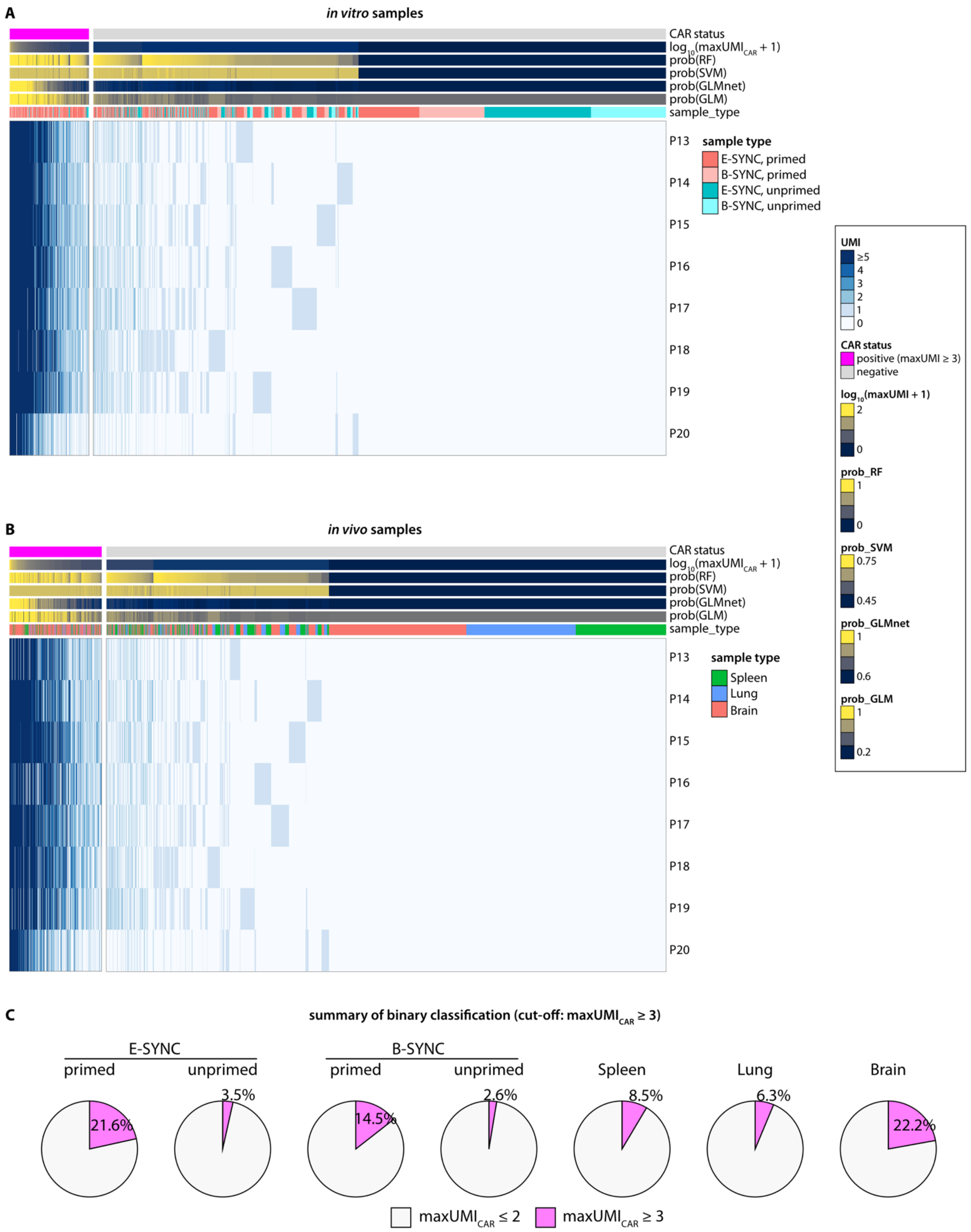

A
in vitro samples
CAR status
log10(maxUMI CAR + 1)
prob(RF)
prob(SVM)
prob(GLMnet)
prob(GLM)
sample_type
P13
P14
P15
P16
P17
P18
P19
P20
sample type
E-SYNC, primed
B-SYNC, primed
E-SYNC, unprimed
B-SYNC, unprimed
UMI
≥5
4
3
2
1
0
CAR status
positive (maxUMI ≥ 3)
negative
log10(maxUMI + 1)
2
0
prob_RF
1
0
prob_SVM
0.75
0.45
prob_GLMnet
1
0.6
prob_GLM
1
0.2
B
in vivo samples
CAR status
log10(maxUMI CAR + 1)
prob(RF)
prob(SVM)
prob(GLMnet)
prob(GLM)
sample_type
P13
P14
P15
P16
P17
P18
P19
P20
sample type
Spleen
Lung
Brain
C
summary of binary classification (cut-off: maxUMI CAR ≥ 3)
E-SYNC
primed
unprimed
B-SYNC
primed
unprimed
Spleen
Lung
Brain
21.6%
3.5%
14.5%
2.6%
8.5%
6.3%
22.2%
maxUMI CAR ≤ 2
maxUMI CAR ≥ 3

**Fig. S14. Classification of CAR-high and CAR-low subsets among synNotch-positive cells.**

(**A–B**) Heatmaps showing the classification of CAR-high and CAR-low cells across *in vitro* (**A**) and *in vivo* (**B**) samples.

(**C**) Pie charts illustrating the proportions of synNotch-positive cells classified as CAR-high in each sample type, defined by a $maxUMI_{CAR} \geq 3$ cutoff.

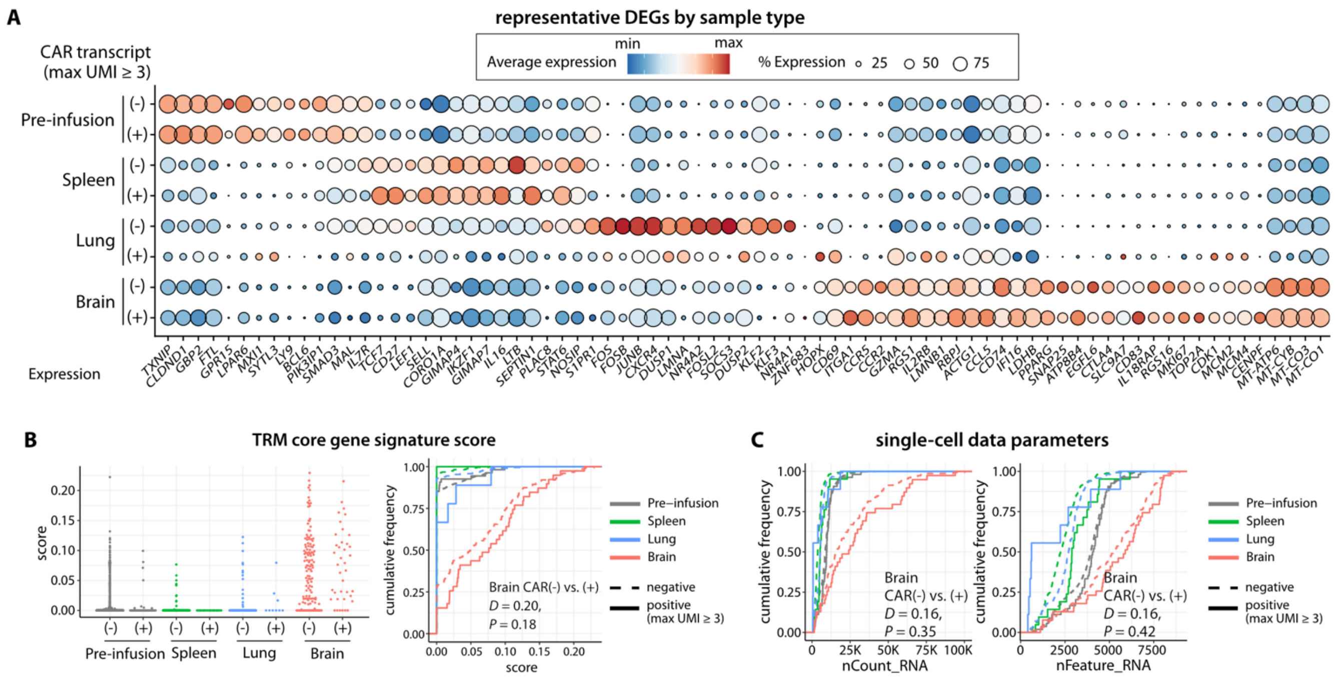


**Fig. S15. CAR transcript abundance and activation state of CD4⁺ synNotch-CAR-T cells (Related to Fig. 7).**

(**A**) Dot plot showing expression of representative DEGs by sample type (same gene list as in **Fig. S9D** and **S12A**), stratified by CAR-expressing ($maxUMI_{CAR} \geq 3$) versus non-expressing cells.

(**B**) Swarm plot and empirical cumulative distribution function (ECDF) plot showing TRM core gene signature score distributions. Differences between CAR-positive and -negative cells within the brain were assessed using the Kolmogorov–Smirnov test (*P* = 0.18).

(**C**) ECDF plot comparing nCount_RNA and nFeature_RNA values between CAR-expressing and non-expressing cells within the CD4⁺ synNotch-positive population. Differences within the brain were assessed using the Kolmogorov–Smirnov test.

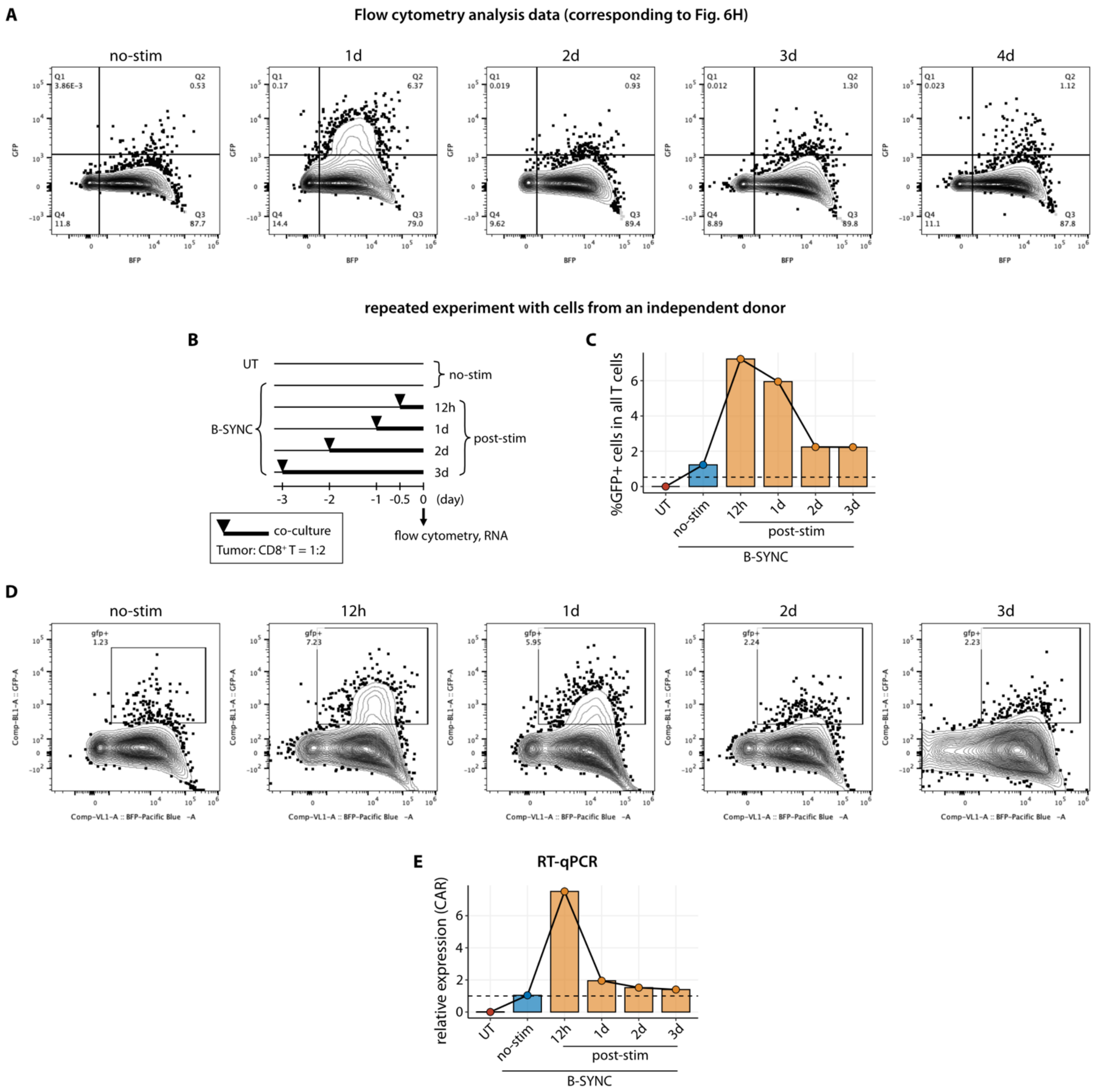


**Fig. S16. Time-course co-culture experiment of B-SYNC cells with GBM6 (Related to Fig. 6).**

(**A**) Flow cytometry contour plots showing cells expressing GFP fused to CAR within the live, singlet, non-tumor (mCherry-negative) population at each time point, corresponding to **Fig. 6H**. (**B**) Schematic overview of an independent repeat of the time-course co-culture experiment of B-SYNC cells with the human GBM cell line GBM6 ($BCAN^+$/$EphA2^+$/$IL13R\alpha2^+$). Co-culture was performed in 24-well plates, with each well containing $0.5\times10^5$ GBM cells and $1\times10^6$ $CD8^+$ B-SYNC cells.

(**C–D**) Flow cytometry analysis showing the percentage of CAR-expressing cells ($GFP^+$) within the live, singlet, non-tumor (mCherry-negative) population (**C**), and the corresponding contour plots (**D**).

(**E**) RT-qPCR data showing CAR transcript abundance across conditions. Relative expression was calculated using the $2^{-\Delta\Delta Cq}$ method, with the B-SYNC no-stimulation (no-stim) condition and the *RPS18* transcript as references.

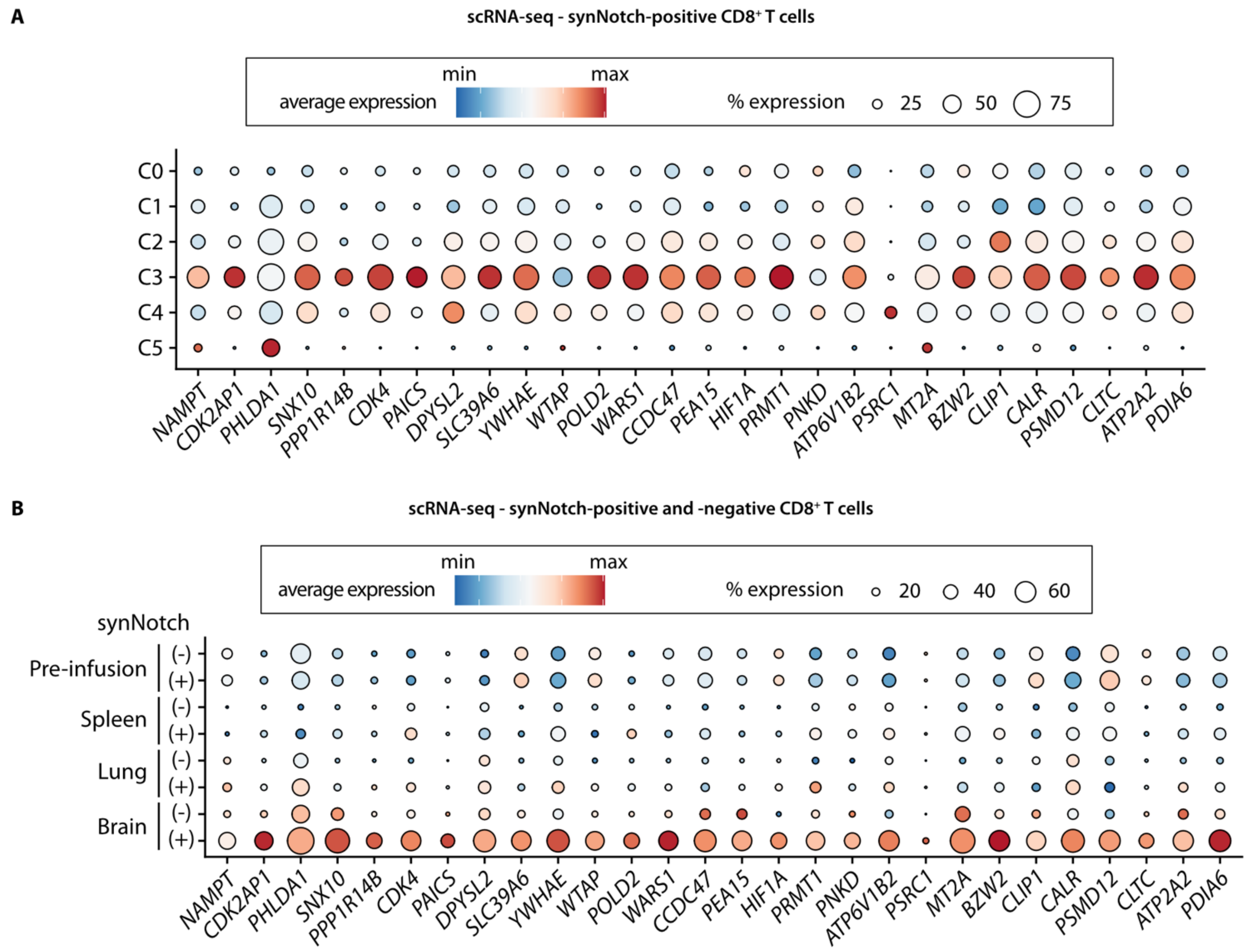


**Fig. S17. Overlap between time-course co-culture bulk RNA-seq and scRNA-seq datasets (Related to Fig. 6L).**

(**A–B**) Dot plots showing expression patterns of representative genes that are upregulated and sustained after co-culture, across CD8+ T cell clusters (**A**) and synNotch-positive and -negative cells from different sample types (**B**) in the scRNA-seq dataset.

### References


1. Y. Chu, E. Dai, Y. Li, G. Han, G. Pei, D. R. Ingram, K. Thakkar, J.-J. Qin, M. Dang, X. Le, C. Hu, Q. Deng, A. Sinjab, P. Gupta, R. Wang, D. Hao, F. Peng, X. Yan, Y. Liu, S. Song, S. Zhang, J. V. Heymach, A. Reuben, Y. Y. Elamin, M. P. Pizzi, Y. Lu, R. Lazcano, J. Hu, M. Li, M. Curran, A. Futreal, A. Maitra, A. A. Jazaeri, J. A. Ajani, C. Swanton, X.-D. Cheng, H. A. Abbas, M. Gillison, K. Bhat, A. J. Lazar, M. Green, K. Litchfield, H. Kadara, C. Yee, L. Wang, Pan-cancer T cell atlas links a cellular stress response state to immunotherapy resistance. *Nat. Med.* **29**, 1550–1562 (2023).

2. M. Andreatta, J. Corria-Osorio, S. Müller, R. Cubas, G. Coukos, S. J. Carmona, Interpretation of T cell states from single-cell transcriptomics data using reference atlases. *Nat. Commun.* **12**, 2965 (2021).

3. M. Andreatta, L. Hérault, P. Gueguen, D. Gfeller, A. J. Berenstein, S. J. Carmona, Semi-supervised integration of single-cell transcriptomics data. *Nat. Commun.* **15**, 872 (2024).

4. L. Zheng, S. Qin, W. Si, A. Wang, B. Xing, R. Gao, X. Ren, L. Wang, X. Wu, J. Zhang, N. Wu, N. Zhang, H. Zheng, H. Ouyang, K. Chen, Z. Bu, X. Hu, J. Ji, Z. Zhang, Pan-cancer single-cell landscape of tumor-infiltrating T cells. *Science* **374**, abe6474 (2021).

5. B. V. Kumar, W. Ma, M. Miron, T. Granot, R. S. Guyer, D. J. Carpenter, T. Senda, X. Sun, S.-H. Ho, H. Lerner, A. L. Friedman, Y. Shen, D. L. Farber, Human tissue-resident memory T cells are defined by core transcriptional and functional signatures in lymphoid and mucosal sites. *Cell Rep.* **20**, 2921–2934 (2017).

6. P. A. Szabo, H. M. Levitin, M. Miron, M. E. Snyder, T. Senda, J. Yuan, Y. L. Cheng, E. C. Bush, P. Dogra, P. Thapa, D. L. Farber, P. A. Sims, Single-cell transcriptomics of human T cells reveals tissue and activation signatures in health and disease. *Nat. Commun.* **10**, 4706 (2019).
